# Building a realistic, scalable memory model with independent engrams using a homeostatic mechanism

**DOI:** 10.1101/2023.08.29.555246

**Authors:** Marvin Kaster, Fabian Czappa, Markus Butz-Ostendorf, Felix Wolf

## Abstract

Memory formation is usually associated with Hebbian learning, using synaptic plasticity to change the synaptic strengths but omitting structural changes. Recent work suggests that structural plasticity can also lead to silent memory engrams, reproducing a conditioned learning paradigm with neuron ensembles. However, this work is limited by its way of synapse formation, enabling the formation of only one memory engram. Overcoming this, our model allows the formation of many engrams simultaneously while retaining high neurophysiological accuracy, e.g., as found in cortical columns. We achieve this by substituting the random synapse formation with the Model of Structural Plasticity (Butz and van Ooyen, 2013). As a homeostatic model, neurons regulate their activity by growing and pruning synaptic elements based on their current activity. Utilizing synapse formation based on the Euclidean distance between the neurons with a scalable algorithm allows us to easily simulate 4 million neurons with 343 memory engrams. These engrams do not interfere with one another by default, yet we can change the simulation parameters to form long-reaching associations. Our model paves the way for simulations addressing further inquiries, ranging from memory chains and hierarchies to complex memory systems comprising areas with different learning mechanisms.

## 1 Introduction

Memory is a key ingredient in our thinking and learning process. However, the understanding of learning and memories is still very limited. In 1921, Richard Semon described the idea of a memory engram, the neurophysiological trace of a memory. Nowadays, working memory is usually characterized by persistent neuron activity (Barak and Tsodyks, 2014). Recent work (Fiebig and Lansner, 2017) modeled this persistent activity with a partly plastic network and synaptic plasticity.

However, synaptic plasticity is limited to changing the weights of already existing connections and prohibits the formation of new synapses between neurons. A memory engram, e.g., in the form of a strongly interconnected group of neurons, could represent an abstract memory. If we now want to form an association with another memory in the form of a memory engram, we are limited to strengthening existing synapses with synaptic plasticity alone. We cannot build an association if there is no or only sparse connectivity between these two engrams. Moreover, ignoring structural plasticity limits the storage capacity of the brain drastically (Chklovskii et al., 2004), and structural plasticity can overcome both problems. An alternative for a model using only synaptic plasticity is to use all-to-all connectivity, but this is impractical for large networks. Additionally, recent work showed that structural plasticity also plays an essential role in the biology of memory formation (May, 2011, Holtmaat and Caroni, 2016, Butz et al., 2009) and especially long-term memory, on the other hand, is usually characterized by silent synapses (Gallinaro et al., 2022).

While most structures (Mizrahi, 2007, Kalisman et al., 2005) and synapses in the developed brain remain stable, there is evidence that learning (Holtmaat and Svoboda, 2009) and sensory input leads to an increased synapse turnover, growth of dendritic spines, and axonal remodeling (Barnes and Finnerty, 2010). For example, the alternate trimming of whisker hair of mice leads to a changed sensory input and an increased dendritic spine turnover (Trachtenberg et al., 2002), with 50% of the formed spines becoming stable. Another example is the change in the brain’s gray matter when adults learn juggling (Boyke et al., 2008). Recently, it was proposed that synaptic plasticity plays a vital role in memory consolidation (Caroni et al., 2012, Holtmaat and Caroni, 2016, Butz et al., 2009). This is the process of transmitting information from the short-term memory into the long-term memory. First, learning increases the synaptic turnover and the formation of vacant synaptic elements (Butz et al., 2009, Holtmaat and Svoboda, 2009, Caroni et al., 2012). Synaptic elements are dendritic spines and axonal boutons, which, if unoccupied, generate potential synapses (Stepanyants et al., 2002). Some potential synapses later materialize, forming actual new synapses (Caroni et al., 2012, Butz et al., 2009). The correlation between spine stabilization and the performance of animals in tests (Xu et al., 2009) supports this hypothesis further. Moreover, when newly formed synapses during training are impaired, the performance of the animal decreases (Hayashi-Takagi et al., 2015), indicating that the formation of new synapses is essential for learning. However, how the learning process interacts with the increased turnover and stabilization of specific spines, without overall modification of the structures in the brain, remains unclear (Caroni et al., 2012).

Dammasch (1990) proposed the idea of forming memory engrams based on structural plasticity. Recently, Gallinaro et al. (2022) developed a model that can form silent memory engrams with only structural plasticity on a homeostatic basis. They showed that a homeostatic rule implicitly models a Hebbian learning rule and demonstrated this principle with a conditioned learning paradigm. However, their model connects neurons uniformly at random, making the simultaneous formation of clustered neurons as distinct memory engrams impossible. The stimulation of multiple memory engrams at once would lead to one single large memory engram instead of multiple small ones, which differs from the brain, where different regions are active simultaneously without interfering with one another. We extended their work with a more neurophysiological accurate approach by connecting neurons depending on their distance and forming memory engrams simultaneously. We simulated 4 million neurons with 343 engrams, enormously increasing the number of simulated neurons and memory engrams simulated with homeostatic structural plasticity. We showed that the engrams do not interfere with each other.

The structure of the model underlying our simulation draws inspiration from the human cortex, which is organized in columns further subdivided into minicolumns (Mountcastle, 1997). Each of these minicolumns consists of excitatory pyramid cells that act as a group for a specific feature depending on the cortex area. For example, a minicolumn in the visual cortex represents the orientation of an object in a visual field (Hubel and Wiesel, 1962). The different minicolumns combined comprise a hypercolumn that enables encoding all orientations for a certain spot on the retina. Similar organizations can be found in the auditory cortex (Reale and Imig, 1980) and the somatosensory cortex (Ruben et al., 2001). The columns of higher associative cortices represent more complex features such as colors (Hadjikhani et al., 1998), objects (Ĺopez- Aranda et al., 2009), or persons (Downing and Peelen, 2011). In our study, we adopted the cortical organization of the associative cortices, where complex features can be learned and represented in a column. Hence, we split our network into boxes that represent a column. Each column represents a feature that acts independently from neighboring columns, and, therefore, all columns can be active simultaneously. In our example with the visual cortex, two neighboring columns encode an orientation for their field of view, respectively, without interference.

Our contributions are:

- We increased the neurophysiological plausibility of homeostatic memory models, which enables us to model cortical-like structures.
- Our model maintains the introduced structure of the cortical columns without additional constraints on how the neurons can connect.
- We formed multiple ensembles in parallel without unwanted interference. This is comparable to the brain with its many concurrent activities.

## 2 Materials and methods

In the experiment, we followed the work of Gallinaro et al. (2022), modeling a conditioned learning paradigm. In a conditioned learning paradigm, a subject learns the relationship between an unconditioned stimulus and a neutral stimulus. Before the experiment, the subject shows an unconditioned reaction to the unconditioned stimulus. During the experiment, both unconditioned (e.g., food) and neutral stimuli (e.g., bell) are presented simultaneously. After the experiment, the subject shows the same previously unconditioned (now conditioned) reaction (e.g., salivating) for the previously neutral (now conditioned) stimulus. At the level of neuronal networks, we modeled a stimulus by stimulating neurons and a reaction as the firing of a downstream, so-called readout, neuron. We split the whole population into boxes of equal side lengths and divided the neurons within each box into four distinct neuron ensembles: Unconditioned stimulus (US), conditioned stimulus 1 (C1), conditioned stimulus 2 (C2), and the rest. We added a single readout neuron (R) to each box that is fully connected to the neurons in the ensemble US with static connections (Fig. 1) to monitor the unconditioned reaction of the network. With these neuron ensembles, we can model a conditioned learning paradigm by representing a neuron ensemble with a stimulus. Like the conditioned learning paradigm, where an unconditioned learning stimulus is presented at the same time as a conditioned stimulus and, therefore, learns a relationship between them, the network shall learn a relationship between US and C1 within each box. However, there should be no relationship associated with C2 or across different boxes, as C2 acts as our control ensemble. To accomplish this, we stimulated the ensembles US and C1 together. We performed these stimulations simultaneously for every memory that we wanted to learn.

**Figure 1:**
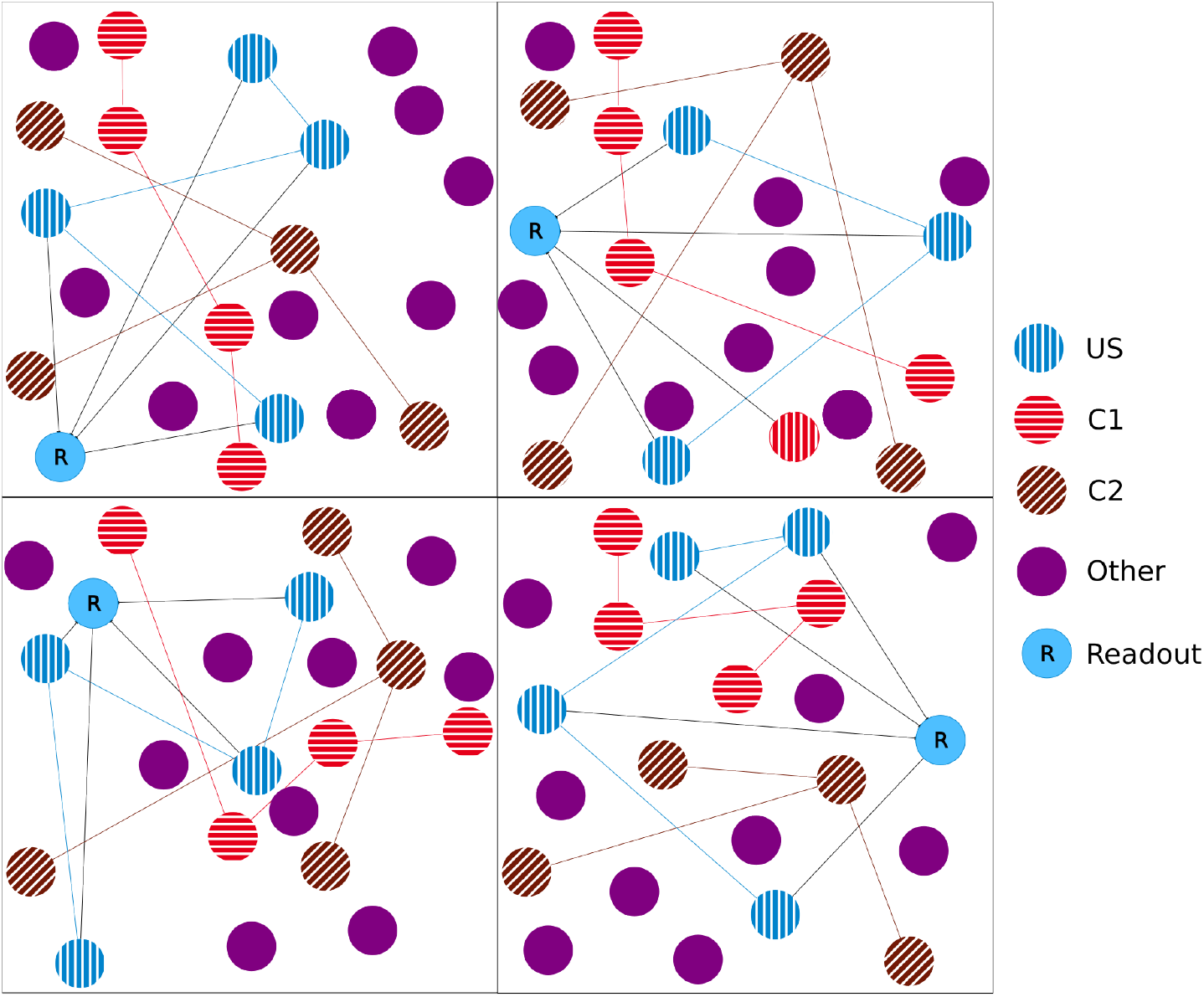
Initial setup of our model network as a 2D simplification with 2×2 boxes. We split the boxes into equally sized boxes and grouped a subset of neurons from each box in three equally sized ensembles representing the unconditional stimulus (US), the first conditional stimulus (C1), and the second conditional stimulus (C2). The neurons of the ensemble US are fully connected with an additional readout neuron (R). The remaining neurons did not belong to any ensemble.

### 2.1 Model of Structural Plasticity

The Model of Structural Plasticity (MSP) (Butz and van Ooyen, 2013) enables structural plasticity based on a homeostatic mechanism. Each neuron controls its excitability and, therefore, the number of synapses based on its calcium concentration, which acts as a proxy for its firing rate. An approximation of MSP (Rinke et al., 2018, Czappa et al., 2023) reduced the model’s computational complexity from *O*(*n*^2^) to *O*(*n* log *n*), which enabled simulations with up to 10^9^ neurons. Using current technology, human-scale simulations with 10^11^ neurons are possible in principle. We provide all model parameters in Tab. 1.

**Table 1:** Parameters for the experiments.

| Parameter | Parameter description | Value |
| --- | --- | --- |
| $\mu_{\text{background}}$ | Mean of the background activity | 5 |
| $\sigma_{\text{background}}$ | Standard deviation of the background activity | 2 |
| $\theta$ | Barnes–Hut acceptance criterion | 0.3 |
| $\tau_{\text{vac}}$ | Retract ratio | 100 |
| $\epsilon$ | Target calcium level | 0.7 |
| $\eta_{\text{axon}}$ | Minimum calcium level to grow axons | 0.4 |
| $\eta_{\text{den\_ex}}$ | Minimum calcium level to grow excitatory dendrites | 0.1 |
| $\eta_{\text{den\_inh}}$ | Minimum calcium level to grow inhibitory dendrites | 0 |
| $\sigma$ | Gaussian scaling parameter | 12 |
| $k$ | Synapse conductance | 3 |
| $\tau$ | Calcium decay constant | 10,000 |
| $\beta$ | Calcium intake | 0.001 |
| $\nu_{\text{axon}}$ | Growth rate of axons | 0.0003 |
| $\nu_{\text{den\_ex}}$ | Growth rate of excitatory dendrites | 0.0006 |
| $\nu_{\text{den\_inh}}$ | Growth rate of inhibitory dendrites | 0.0006 |
| $a_{\text{izh}}$ | Izhikevich model parameter | 0.1 |
| $b_{\text{izh}}$ | Izhikevich model parameter | 0.2 |
| $c_{\text{izh}}$ | Izhikevich model parameter | -65 |
| $d_{\text{izh}}$ | Izhikevich model parameter | 2 |

#### 2.1.1 Growth model

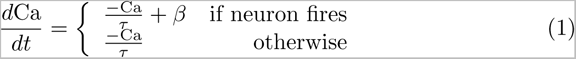

The calcium concentration that acts as a proxy for the firing rate is described in Eq. 1. It decreases over time with the time constant *τ* and increases by *β* when the neuron fires. Based on the calcium concentration, the number of synaptic elements of a neuron grows or shrinks as described in Eq. 2. It depends on the growth rate *ν*, the current calcium concentration Ca, the minimum calcium concentration *η* required to grow synaptic elements, and the target calcium rate *ɛ*. Hence, if the calcium concentration is higher than the target, the neuron prunes synaptic elements; it grows new elements if the concentration is lower than the target. We follow Butz and van Ooyen (2013) for selecting our minimum calcium concentration 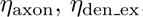, and 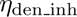, where they showed that a high *η*_axon_ and a low *η*_den_ matches the experimental data and enables functional remapping.

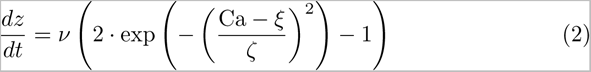

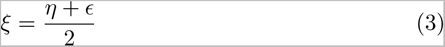

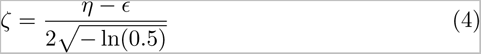

#### 2.1.2 Forming and pruning of synapses

Since we already determined how many synaptic elements a neuron forms, we must now actually form synapses between unbound synaptic elements or prune them if the number of grown synaptic elements of a neuron is smaller than its actual synapses. In the last case, we chose a synapse to prune uniformly at random from all synapses of the neuron. Consequently, the synapse will be removed from the source and target neuron, independently of the number of grown synaptic elements of the partner neuron. When we need to form a new synapse, we search for vacant dendritic elements for every vacant axonal element. We choose a target element randomly based on weighted probabilities. Each target dendritic candidate is weighted with a probability based on the distance between the positions *x_i_* of the neuron of the inititaiting axonal element and the candidate’s neuron position *x_j_*, as described in Eq. 5. Note that the implementation uses an approximation (Rinke et al., 2018, Czappa et al., 2023) so that it does not need to calculate the probability for all candidates.

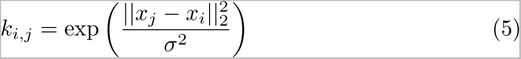

Synaptic elements that are not connected are removed over time depending on the time constant *τ*_vac_, as described in Eq. 6.

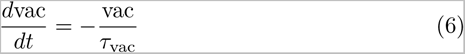

#### 2.1.3 Electrical activity

The electrical input *I* is the sum of three inputs: synaptic, background, and stimulation, as described in Eq. 7. The synaptic input (Eq. 8) is calculated over all fired input neurons *j*. Each of these synapses has a weight that of 1 for excitatory input neurons or *−*1 for inhibitory ones and is multiplied by the fixed synapse conductance *k*. The sum over those products for all input synapses of neuron *j* whose source neurons fired is the synaptic input for *j*. Additionally, the neurons are driven by a random, normally distributed background activity (Eq. 9).

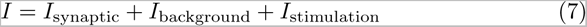

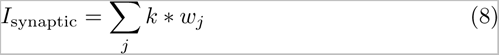

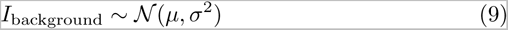

### 2.2 Izhikevich model

We chose the Izhikevich model (Izhikevich, 2004) as our neuron model because it enables modeling different spiking patterns and strikes a good compromise between biological accuracy and computational efficiency. We describe the membrane potential in Eq. 10 where *k*_0_, *k*_1_, and *k*_2_ are constants and *u* is the membrane recovery variable as described in Eq. 11. The membrane recovery depends on the fixed parameters *a*, *b*, and *c* and accounts for the hyperpolarization period of a neuron. Finally, we assumed that a neuron spikes if the membrane potential reaches 30 mV. Then, we reset the membrane potential and membrane recovery variables according to Eq. 12.

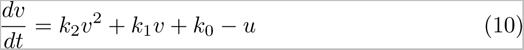

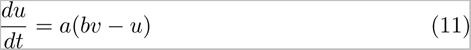

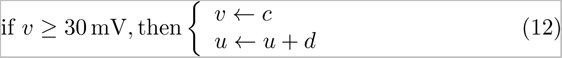

### 2.3 Network setup and stimulation

Unless noted otherwise, our network consisted of 337,500 neurons, each with 20% inhibitory and 80% excitatory neurons. We distributed the neurons uniformly at random in a 3-dimensional cubic space. We split the network into 3 × 3 × 3 = 27 equally sized cubes with about 12,500 neurons each, following Gallinaro et al. (2022). In each box, we grouped a part of the neurons into disjoint ensembles of 333 excitatory neurons. We selected them randomly from all neurons in the box. Fig. 1 visualizes the setup as a 2D example.

We started our network in a fully disconnected state and let it build up the connectivity from scratch for 1,000,000 steps. In more detail, the neurons received a normally distributed background activity such that they have a certain baseline activity. As this baseline is insufficient to reach the neurons’ target calcium level, they grow synapses to connect to other neurons and receive more input. When the neurons reach the target calcium level, they stop growing synapses and they perform only small modifications to their synapses due to the fluctuations in the input noise and the resulting fluctuations in the neurons’ firing rate. In this state, the network is in equilibrium, as every neuron is approximately at its target calcium level, and only small modifications are made to the network. We saved the newly formed network and used it as a starting point for our experiment. Hence, we continued the simulation with this network and waited for 150,000 steps until the network stabilized itself. Then, we continued the simulation with three stimulation phases similar to Gallinaro et al. (2022): baseline, encoding, and retrieval. We performed all stimulations for 2,000 steps with 20 mV. In the baseline phase, we stimulated all US ensembles in step 150,000, followed by all C1 ensembles (250,000), and lastly, all ensemble C2 (350,000). Note that US, C1, and C2 were stimulated after each other, but all ensembles of the same type across all boxes were stimulated at the same time. The goal of this phase is to strengthen the connectivity within each neuron ensemble. We selected the pause between the stimulation so that the neurons’ calcium levels return to their target level to ensure that there is no priming effect. Moreover, the C2 ensemble acts as a control to counter a priming effect as well as we would see an effect not only in US and C1 but also in C2.

Then, in the encoding phase, we wanted to learn the relationship between US and C1 and, therefore, stimulated all US and C1 ensembles together in step 450,000 and all C2 ensembles in step 550,000. The stimulation of the C2 ensembles alone acts as a control against a priming effect. That would be the case when US or C1 form connections to C2 despite it being stimulated alone. Similar to the baseline phase, we stimulated the specified ensemble types from each box at the same time.

Later, in the retrieval phase, we turned the plasticity off so that no further modifications were possible. We did this to see how the network behaves without any further modifications. Here, we stimulated, beginning in step 650,000, all C1 ensembles consecutively in isolation, one after the other, with a pause of 18,000 steps between them. Note that this is different from the previous two phases, where we stimulated the same ensemble type (e.g., US) from each box at the same time. In the end, we stimulated all C2 ensembles together at step 1,190,000 as a control. We expect that the readout neuron of a box fires at an increased rate when we stimulate C1 of the same box due to the learned relationship and not if we stimulate the C2 ensembles. The protocol is visualized in Fig. 2a.

**Figure 2:**
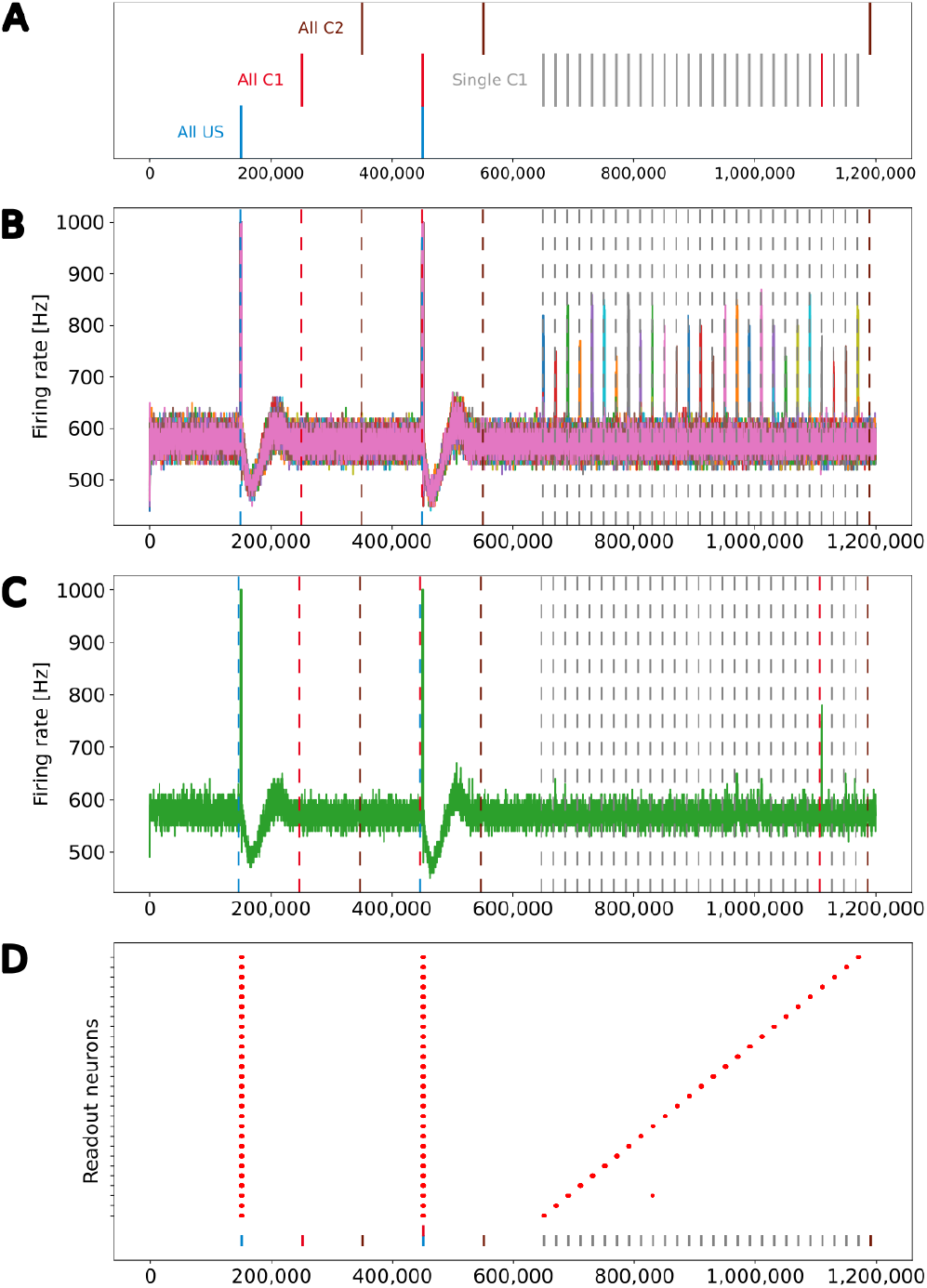
Stimulation protocol and firing rates (y-axis) of all readout neurons plotted over time (x-axis). The vertical dashed lines represent the stimulation of the ensembles US (blue), C1 (red), and C2 (brown). The gray lines and red line on the right represent single stimulations of C1. The readout neurons fire at an increased rate when US is stimulated in the baseline and encoding phase. If the model successfully learns the relationship between the ensembles US and C1 of the same box, neurons of ensemble US will also fire at an increased rate. **(A)** Stimulation protocol of the experiment. The experiment is split into three stimulation phases: baseline, encoding, and retrieval. **(B)** Firing rates of all 27 readout neurons laid over each other. **(C)** Firing rate of a single readout neuron. **(D)** Scatter plot when readout neurons fire at an increased rate. The 27 readout neurons are distributed along the y-axis. A red dot indicates that a readout neuron fires at an increased rate. The stimulation of all US (blue), all C1 (red), all C2 (brown), and single C1 ensembles (gray) are marked with a vertical bar at the bottom.

### 2.4 Validity check

To check the validity of our results, we needed to make an assumption about the distribution of the neurons’ firing rate. We hypothesized that the firing rates follow a normal distribution and showed this with the Kolmogorov–Smirnov test. It calculates the probability that the distribution of our data is not the same as the given distribution to which we compare (in our case: normal distribution).

For this, we simulated the network that resulted from our main experiment for 100,000 steps and recorded the times the readout neurons fire. Then, we split the steps into bins with the size of 1,000 steps and calculated the firing rate of each readout neuron for each bin. As we hypothesized a normal distribution, we calculated the mean and standard deviation for the calculated firing rates. The histogram of the firing rates is shown in Fig. S1. We applied the Kolmogorov– Smirnov test (Massey Jr, 1951) to the calculated firing rates and retrieved a *p* value of 5.34 *∗* 10*^−^*^6^. As a value of *p* smaller than 0.05 can be interpreted as non-significant, we can assume that our firing rates are normally distributed.

Now, we can continue with our validity check by checking the firing rate of the readout neuron during the retrieval phase to ensure that the network behaves as expected. We assumed that the firing rates follow a normal distribution and detect abnormal behavior of the readout neuron when its firing rate is significantly outside of the normal distribution. For it, we used the 3-*σ* rule (Pukelsheim, 1994), as described in Def. 13 with *f* as the firing rate of the readout neuron, *µ* as its mean, and *σ* its standard deviation without any stimulation. 99.7% of the normal firing rate lies within the interval of three times the standard deviation around the mean of the normal distribution of the firing rate. We considered a firing rate outside of this interval for at least 500 steps to be significantly different from the normal firing rate of the neuron when the network is in an uninfluenced state.

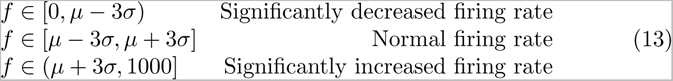

We expected a readout neuron to fire at an increased rate if

- We stimulate its ensemble US during the baseline phase or
- We stimulate its ensembles US and C1 during the encoding phase, or
- We stimulate its ensemble C1 during the retrieval phase It should not fire at an increased firing rate otherwise.

We allowed the increased firing rates to start (end) within 100 ms of the start (end) of the associated stimulation to allow for some variation in the neurons’ behavior.

### 2.5 Large-scale formation of memory engrams

We successfully repeated the experiment with a larger network to show how well our approach scales. Instead of 3 *×* 3 *×* 3 = 27 boxes, we split a larger network into 7 *×* 7 *×* 7 = 343 boxes with a total of 343 *·* 12, 500 = 4, 287, 500 neurons and stimulated the network following the same pattern as before. We analyzed the firing rates of the readout neurons to check whether they fired only when US was stimulated during the baseline and encoding phases and only when C1 in the same box was stimulated during the retrieval phase as described in Sec. 2.4.

### 2.6 Advanced simulations

The following two experiments started with the network that was stimulated as described in Section 2.3 because they analyze the newly created engrams.

#### 2.6.1 Pattern completion

Gallinaro et al. (2022) showed that the stimulation of 50% of the neurons of a single engram was necessary to increase the activity of the rest of the engram; we will improve upon that in both the necessary threshold and the technical contribution. While we stimulated all neurons of ensemble C1 during the retrieval in the previous experiment, we now show that it is sufficient to stimulate only a subgroup of neurons in ensemble C1 to trigger the readout neuron in the associated ensemble US. Hence, after the previous learning experiment with 27 boxes, we continued simulating the network, picked a single ensemble C1 from one box, and stimulated varying numbers of neurons. To this end, we continued the simulation and waited for 60,000 steps until the network stabilized. Then, we turned the plasticity off and stimulated 5% randomly chosen neurons of the ensemble C1 of the center box for 2,000 steps. After a pause of 98,000 steps, we added a further 5% randomly chosen neurons of the ensemble to the stimulated neurons and stimulated them again. We repeated this cycle until we stimulated 100% of the ensemble.

#### 2.6.2 Forming long distance connections

During our main experiment, we wanted to build memory engrams only within a box. Now, we want to show that we can still form connections over long distances and therefore combine memory engrams from different regions. For this, we continued the simulation with a larger Gaussian scaling parameter *σ*_distant_ that makes connections over long distances more probable. We selected two edge boxes on the opposite side of the simulation area to maximize the distance. We waited for 150,000 steps so that the network could stabilize first. Then, we stimulated the ensembles US and C1 from both boxes together for 2,000 steps. After a pause, we turned the plasticity off in step 250,000 and stimulated each ensemble C1 from the boxes after each other with a pause of 8,000 steps between them to check if the network learned the relationship between the ensembles in the two boxes without interfering with other boxes. As a last stimulation in step 520,000, all of the ensembles C2 were stimulated to check the response to the control ensembles.

### 2.7 Ablation studies

We investigated how the network reacts to applied lesions. We are interested in two cases: The loss of synapses and the loss of neurons. For both cases, we continued the simulation from our main experiment and applied the lesion after 150,000 steps. Then, we waited 200,000 steps for the network to stabilize again so that each neuron reached its calcium level again. In step 350,000, we turned the plasticity off and stimulated all C1 ensembles after each other and all C2 ensembles as described for the retrieval phase in Sec. 2.3. We randomly selected a center of the lesion in each box and selected a fraction of neurons that are closest to this center. In step 150,000, we removed all connections from and to the selected neurons. If we decided to lesion the neurons, we also removed them from the network so they were not available for reconnecting their synapses.

## 3 Results

This section is divided into multiple parts. We will start with the analysis of the process of the formation of a single memory engram. Then, we will discuss the simultaneous formation of multiple memory engrams. Finally, we provide the results of the large-scale and advanced simulations.

### 3.1 Process of engram formation

Our experiment consists of three phases: baseline, encoding, and retrieval. We started with the baseline phase. As described in Sec. 2.3, the network was in a stable state with only minor changes in the connectivity at the beginning of the baseline phase (Fig. 3). At this point, the network’s connectivity was random based on the neurons’ distances (Fig. 4b). This phase aims to form neuron ensembles with strong connectivity within the ensemble. The additional stimulations resulted in an increased firing rate (Fig. 3a) of the neurons, which in consequence lead to an increase in their calcium level and therefore to a pruning of synapses (Fig. 3b). After the stimulation ends, the firing rate fell before its usual level due to the smaller synaptic input caused by the pruning of synapses. Consequently, the neurons within the ensemble started regrowing their synapses until they reached a stable firing rate and calcium level as its target value. All neurons within the ensemble regrew their synapses simultaneously and are, therefore, looking for new synapse partners at the same time. This high ratio of potential partners within the same ensemble results in strong connectivity within this ensemble (Fig. 3c). We stimulated a single ensemble, such as US, from each box at the same time. This and the property of forming new synapses based on the distance of neurons results in a strong within-connectivity of the ensemble in a box. After a short delay, we continued with the second C1 ensembles, and after another delay with C2 ensembles. As a result, we had three strongly within-connected ensembles in each box representing an engram (Fig. 4c). We refer to the work of Gallinaro et al. (2022) for details of this process. Fig. 4 shows the process of reorganizing the connectivity during the entire experiment.

**Figure 3:**
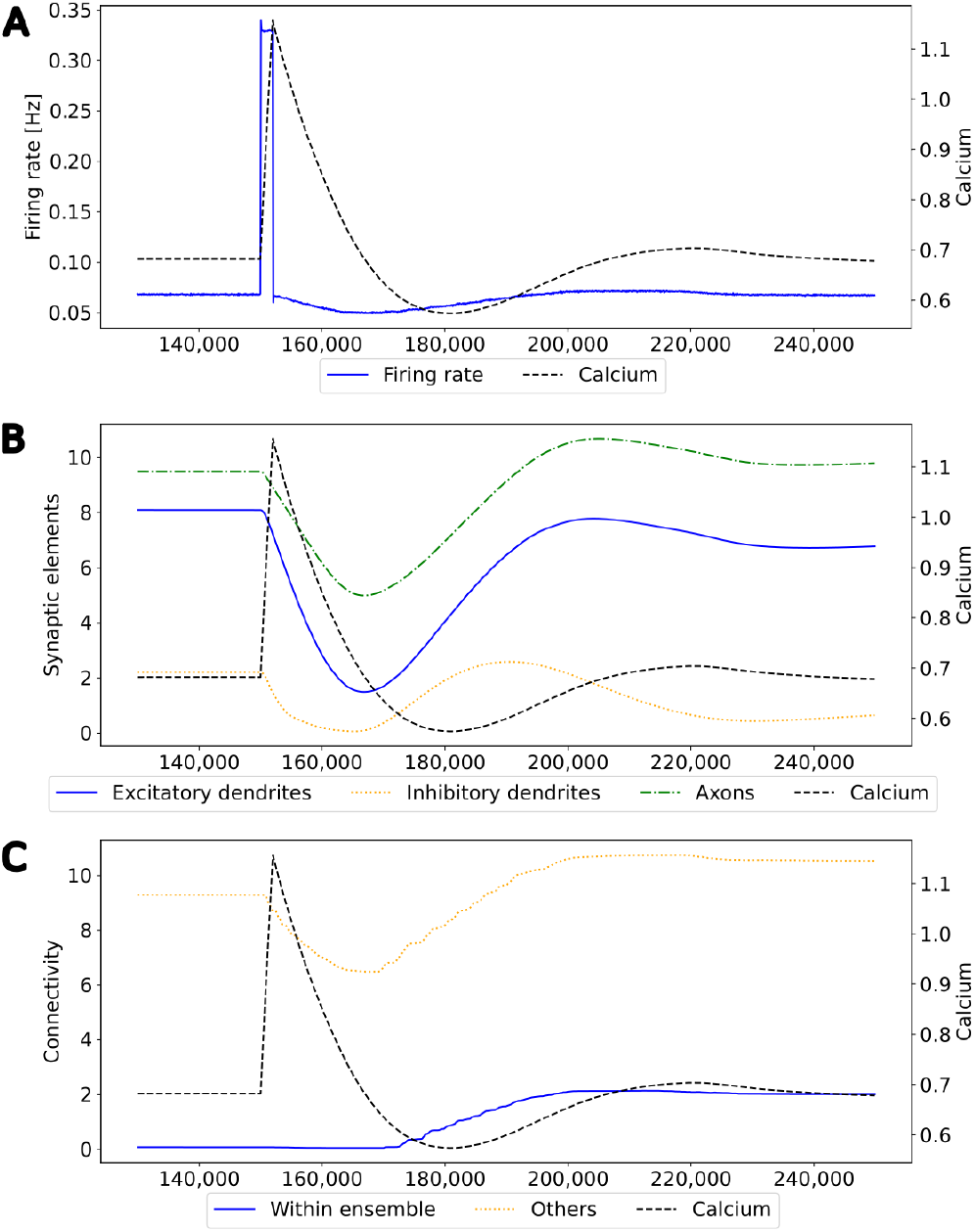
Mechanism of a single engram formation shown with a single ensemble group US. The stimulation in step 150,000 starts the engram formation. **(A)** Intracellular Calcium (right y-axis) as an indicator for the average firing rate (left y-axis) of all neurons in a randomly selected example US ensemble. The stimulation is visible as a high spike in the firing rate. This was followed by decreased activity caused by homeostatic network reorganization as depicted in panels B-C. **(B)** Changes in intracellular calcium triggered growth of synaptic elements of the neurons in the exemplary ensemble. The synaptic elements decrease after the stimulation as calcium was higher than the homeostatic setpoint and grew again afterwards when activities fell below the set-point as a consequence of compensatory pruning of synapses. **(C)** Average connectivity of an exemplary ensemble to itself and other neurons. As a consequence of the changing number of synaptic elements, we see a drop in the connectivity to neurons outside of the ensemble directly after the stimulation which was slowly restored afterwards. Connectivity within the ensemble increased likewise.

**Figure 4:**
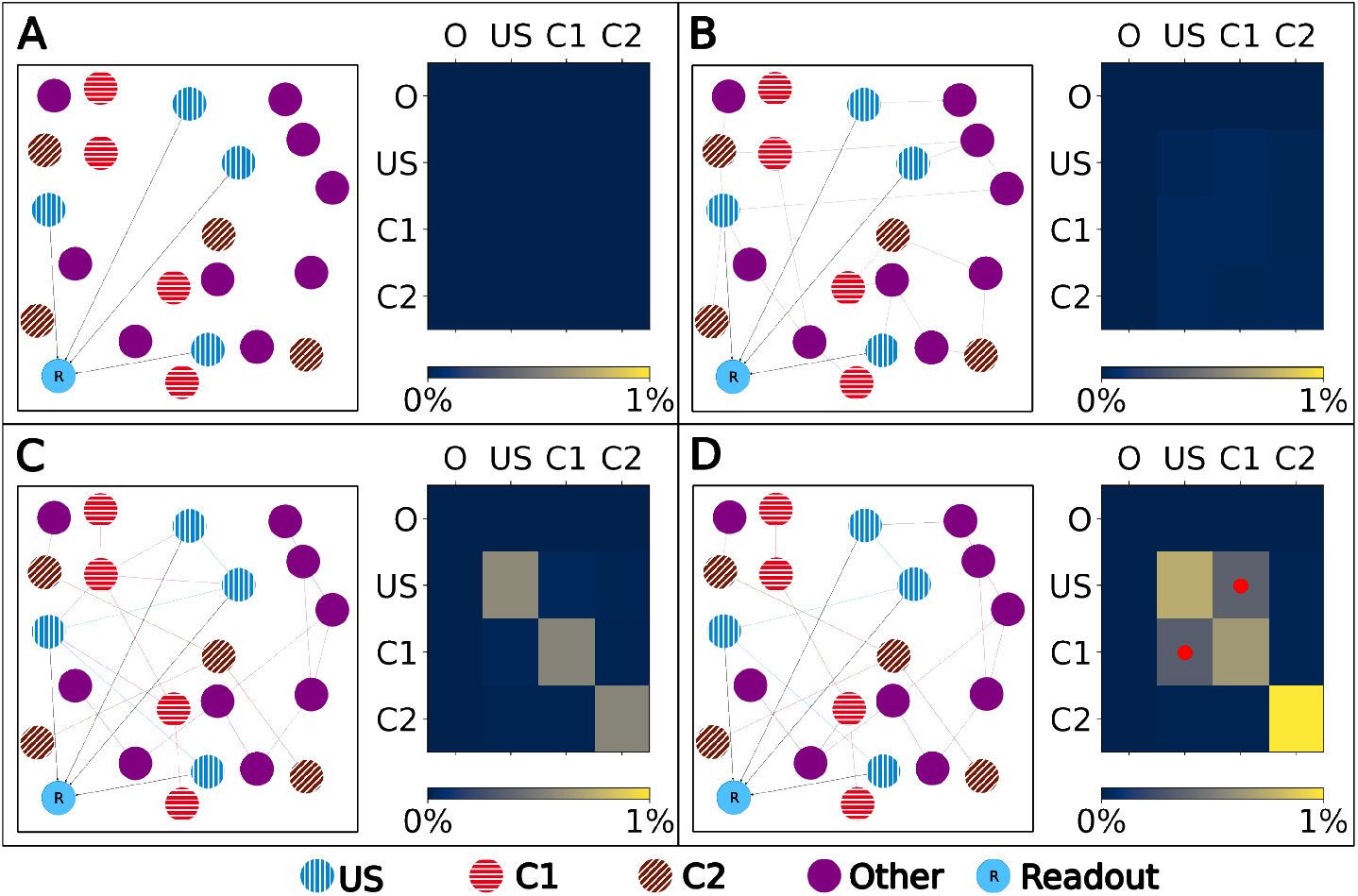
Left: Schematic illustration of the connectivity within a single box. Right: Heatmap of the neurons’ normalized number of input (row) and output (column) connections of each ensemble (O), US, C1 and C2. Normalized with the size of the source and target ensemble.. **(A)** Visualization of the network at the beginning with no connectivity besides the static connections to the readout neuron. **(B)** Visualization of the network after the growth period with random connectivity. **(C)** Visualization of the network after the baseline phase with increased connectivity within each ensemble. **(D)** Visualization of the network after the encoding phase showing that the learned relationship between US and C1 is brought about by an additional increased connectivity between the ensembles C1 and US as marked in the heatmap with red dots.

Next, we continued with the encoding phase. In this phase, the model learned the association between the memory ensembles US and C1 within each box. For this, we stimulated the ensembles US and C1 together. Consequently, the ensembles US and C1 in a single box form many connections using the same mechanism as in the baseline phase (Fig. 4d).

Finally, in the retrieval phase, we checked if the model learned the association between US and C1. The firing rate of an exemplary single box’s readout neuron is visualized in Fig. 2c. Besides the two spikes during the earlier phases when US was stimulated, the readout neuron fired only at an increased rate when the ensemble C1 from the same box was stimulated and not if we stimulated our control ensemble C2 or any other ensemble from another box.

### 3.2 Simultaneous formation of memory engrams

Our network consisted of 3 *×* 3 *×* 3 = 27 boxes, each with three ensembles: US, C1, and C2. By plotting the readout neurons in Fig. 2b and 2d, we show that the network learns the relationships between US and C1 in each box and not between US and C2. Without stimulation, the readout neurons fired at about 600 Hz. All readout neurons fired at steps 150,000 and 450,000 with the maximal frequency of 1000 Hz. This was followed by a decrease of the firing rate to about 450 Hz until it recovers to its normal frequency of 600 Hz. In step 150,000, we stimulated the US ensembles directly connected to their readout neurons during the baseline phase. During the stimulations of C1 in step 250,000 and C2 in step 350,000, the readout neurons did not fire at an increased rate compared to their baseline frequency. This shows that the network works as desired because the unconditioned reaction is shown after the unconditioned stimulus. Later, during the encoding phase, the readout neurons fired again, as expected when US and C1 were stimulated together in step 450,000 but not when C2 was stimulated in step 550,000. Finally, in the retrieval phase, we stimulated the ensemble C1 from each box after each other. The readout neuron of the same box fired when its ensemble C1 was stimulated, indicating that the network learned the relationship between US and C1 only within the same box but not beyond box boundaries. The firing rates were lower than during the stimulation in the encoding phase but still very distinguishable from the rest of the activity. We noticed small spikes of some readout neurons when another box was stimulated. However, the stimulation of our control ensembles C2 still did not influence the readout neurons.

The learned relationship between US and C1 in each box can be explained with the newly formed connections, as visualized in Fig. 5. At the beginning of the simulation, all ensembles were randomly connected (Fig. 5a). After the baseline phase, the connectivity within the ensembles increased, and the connectivity between different ensembles decreased (Fig. 5b). This increased connectivity within ensembles indicates the formation of memory engrams similar to Gallinaro et al. (2022). However, in contrast to what they did, we stimulated all US and C1 ensembles together only once instead of thrice. After the encoding phase, the connectivity between the ensembles US and C1 in the same box increases significantly, clarifying that the network learned the relationship between US and C1 (Fig. 5c). However, we could observe a slightly increased connectivity between the ensembles of neighboring boxes due to the connectivity probability being based on the distance of neurons. This explains the slightly increased activity of the readout neurons sometimes when another box is stimulated. Furthermore, the C2 ensembles formed connections primarly within their ensembles and did not connect to neurons in other boxes or ensembles, confirming that these control ensembles did not influence the learning process.

**Figure 5:**
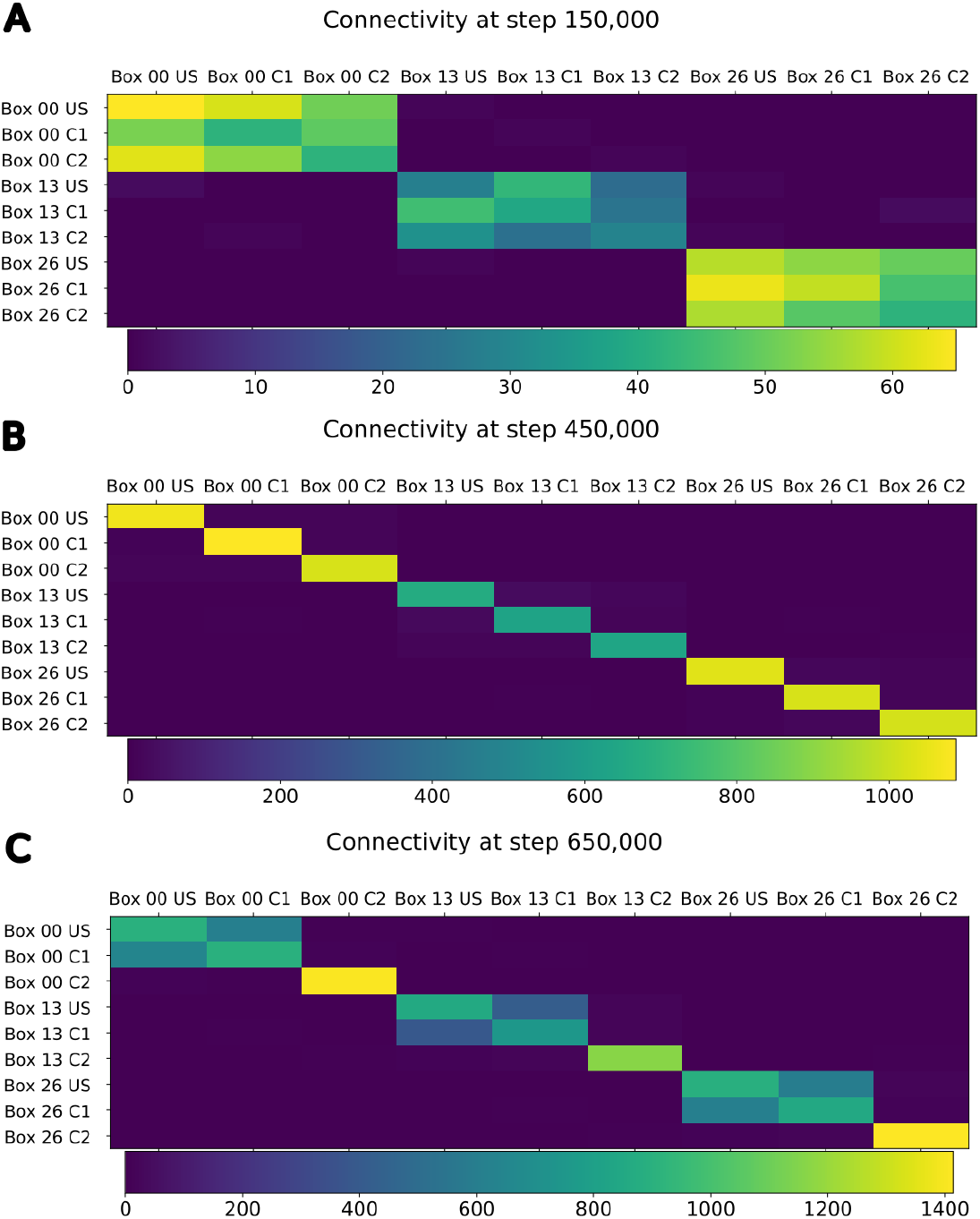
Connectivity between chosen ensembles in the network. From the top left box of the first layer (box 00) over the center box of the second layer (box 13) to the bottom right box of the last layer (box 26). **(A)** During baseline phase, before first stimulation at step 150,000, connectivity is randomly distributed but predominantly remains within a box. **(B)** After baseline phase at step 450,000, connectivity has increased within each neuron ensembles. **(C)** After the encoding phase at step 650,000, connectivity also increased between ensembles US and C1 within the same box.

### 3.3 Spatiotemporal dynamics of homeostatic engram formation

We observe the average calcium level, grown axons and excitatory dendrites, and the connectivity of the neurons over the entire reorganization process for the ensembles US, C1, and C2 for an exemplary single box in Fig. 6 to analyze the mechanism behind the synaptic reorganization further. Moreover, we visualize the average calcium level and, therefore, indirectly, the activity of the neuron groups at different times in the context of the growth curves for the different types of synaptic elements in Fig. 7a. Before the first stimulation in the encoding phase of the ensembles US and C1, the calcium level of all groups was around their target level (Fig. 7a left). The values of the growth curves for this activity level were at around zero as the model is in an equilibrium state with only minor modifications to the network, which is also visible in the almost constant number of axons and dendrites, as illustrated in Fig. 6c,d. The connectivity between US and C1 was almost zero, as there is no relationship between them yet. Instead, the connectivity from within the ensemble US to external neurons was only high for neurons outside of any ensemble just because of the higher number of neurons not belonging to an ensemble compared to the size of the ensembles C1 and C2. Next, we applied the stimulation to the US and C1 ensembles (Fig. 6 (0)), causing their activity to increase and moving the calcium for these ensembles to the right in Fig. 7a (center). The values of the ensembles’ growth curves were almost −1 for this calcium level, reducing the synaptic elements at maximum speed. Consequently, the number of axons and dendrites and their connectivity started decreasing.

**Figure 6:**
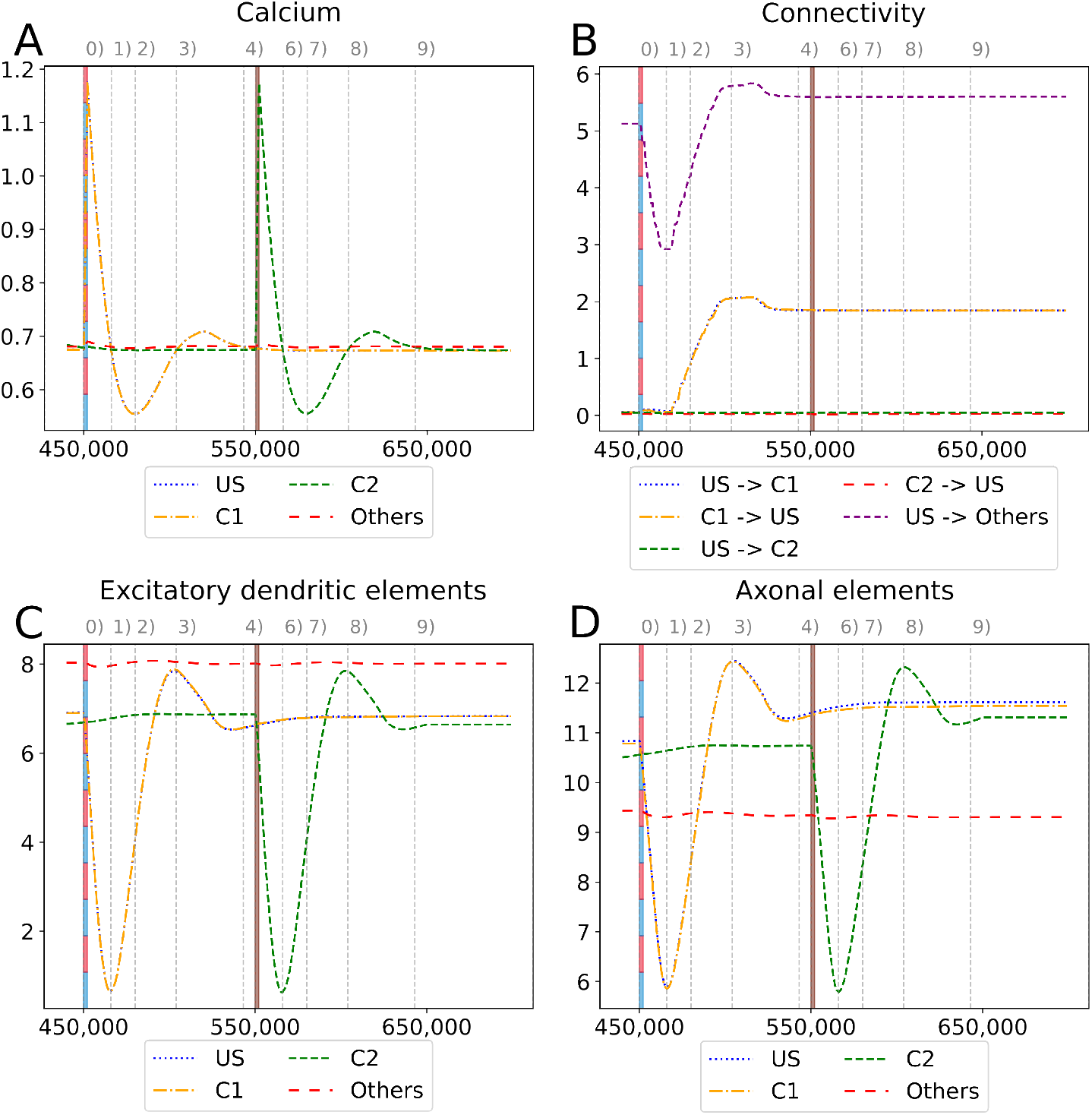
Homeostatic reorganization of neuron ensembles within a single box during the encoding phase. All graphs are averaged over all neurons of the respective ensemble within a single box. We stimulated the ensembles US and C1 at step 450,000 (vertical red-blue line) and the ensemble C2 at step 550,000 (vertical brown line). Panels show the average intracellular calcium concentration **(A)**, connectivity **(B)**, amounts of excitatory dendritic elements **(C)** and axonal elements **(D)**. Networks are in a homeostatic equilibrium before first stimulation **(0)**. After stimulation, activity of the ensembles US and C1 is increased, resulting in pruning of synapses until calcium levels fall below the target value **(1)**. Then, synaptic elements are formed, and calcium level may rise again **(2)**. Axonal and dendritic elements are simultaneously formed by neurons of ensembles US and C1, which become available for synapse formation and explain the observed connectivity increase between these ensembles. The growth phase is followed by a transient overshoot **(3)** and subsequent minor pruning until a homeostatic equilibirum is reached again **(4)**. The stimulation of C2 **(5)** follows the same trend (**(5)**-**(9)**) with the main difference that neurons from ensemble C2 are the only ones that grow axonal and dendritic elements at the same time. The consequence is a massive increase of connections within the ensemble but not between C2 and ensemble US.

**Figure 7:**
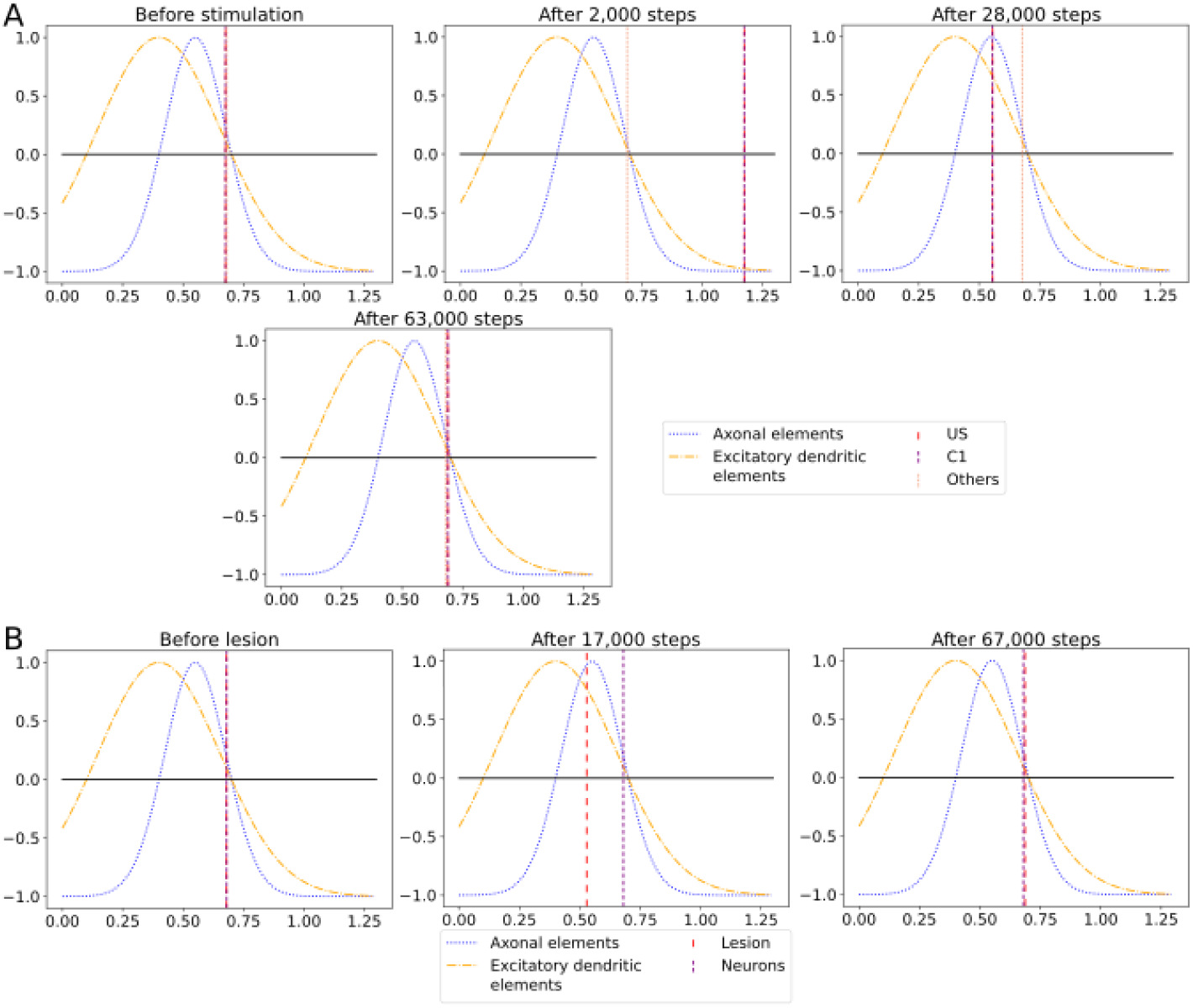
The average calcium level of groups of neurons of a single box (x-axis) with calcium-dependent growth curves (y-axis). **(A)** Calcium level before and after the ensembles US and C1 were stimulated at step 450,000. Subsequentially, their calcium levels increased and growth curves caused synaptic elements to decrease and synapses to prune. When stimulation stopped and synapses were pruned, activities and calcium levels, respectively, dropped below the homeostatic set-point which, in turn, triggered the growth of synaptic elements and potentially also the growth of new synapses. Note, that calcium levels below the set-point were in an optimal regime for axonal element formation while dendritic elements grew slower. A surplus of axonal elements may result in more long-range connections while a prolongued growth of dendritic elements extended the phase in which new engrams could form because it may take longer until activities return to a homeostatic set-point. **(B)** Average calcium levels during ablation studies in which we removed connectivity of 50% of the neurons in a box. Directly after the stimulation, calcium levels of lesioned neurons dropped due to the lack of input. As a result, the neurons start regrowing synaptic elements until enough synapses were formed to restore activity homeostasis. The homeostatic reorganization is comparable to engram formation after stimulation in A. Note, that even for higher deletion rates neurons will return average firing rates to the homeostatic set-point (data not shown) very much as in B, however without functional recovery of trained engrams.

After the stimulation ended, the calcium levels fell below their target (Fig. 6 (1)). The neurons started rebuilding their synapses as soon as the calcium level dropped below its target level. The decrease in the calcium level was slowed down until it reached its lowest level (Fig. 7a right, Fig. 6 (2)). At this calcium level, the ensembles US and C1 correspond to the spike of the growth curve of the axons. The growth curves of the dendrites are lower but still positive. Hence, the ensembles US and C1 build new synapses fast, as shown in Fig. 6c,d. As the ensembles US and C1 built new synapses simultaneously, they formed their synapses, to a large degree between themselves, increasing connectivity between them (Fig. 6b). The fast formation of synapses caused the calcium level to exceed its target (Fig. 6 (3)) due to the increased synaptic input and, therefore, led again to a slight reduction of synaptic elements. In the end, the calcium levels of all groups returned to their set level (Fig. 7a bottom, 6 (4)) and the connectivity as well as the number of synaptic elements remained at a stable level. During the entire stimulation and reorganization of the ensembles US and C1, the calcium level, number of synaptic elements, and connectivity of the control ensemble C2 remained unchanged. The reaction of the ensemble C2 followed the same trend as the ensembles US and C1 during and after its stimulation (Fig. 6 (5)-(9)). As we stimulate the ensemble C2 on its own, we see no effect on the other ensembles.

When we look at the course of the calcium levels before (Fig. 7b left) and after (Fig. 7b center) we removed all connectivity between a group of neurons (Fig. 7b), we observe that the levels dropped from their target level for the lesioned group of neurons. The decreased calcium level corresponds almost to the peak of the growth curve for axons (Fig. 7b center), meaning that axons were built at their maximum speed. Moreover, dendrites were also built fast but not at their maximum speed. As a result of the build-up of synaptic elements, the activity and, therefore, the calcium levels of the lesioned neurons started increasing until they reached their target level again (Fig. 7b right). The calcium level of the non-lesioned neurons remained mostly unchanged. The rebuilding of the synaptic elements after completely removing them for the lesioned neurons is similar to rebuilding the synapses after the stimulation during the conditioned learning experiment. This explains why the network can only recover its learned relationships if all or a high ratio of the lesioned neurons are part of the learned relationship. Otherwise, the network expresses the same learning effect as before but between all lesioned neurons regardless of the affiliation of the neurons.

As we could observe, the synaptic reorganization in our model always follows the same pattern. First, there is a loss of connectivity either directly caused by a lesion or indirectly caused by stimulation and the following pruning of synaptic elements caused by the increased activity and the growth rule of our model itself (Fig. 6 (0)-(1)). In consequence, the activity of the neurons decreases because of the smaller synaptic input. Then, neurons start rebuilding the synapses, connecting mostly among themselves (Fig. 6 (1)-(4)). In the end, the activity of the neurons returns to its initial level. This contrasts Hebbian plasticity, where we cannot observe such a sequence of events. Neurons with similar activity patterns, e.g., because of stimulation, directly strengthen their connections. Strengthening their connections makes it more likely that they fire together and increase their activity. As they are now more likely to fire together, Hebbian plasticity continues to strengthen their synapses. This can lead to continuous strengthening of synapses and increasing neuron activities with unbound synaptic weights. Multiple counter mechanisms have been discussed (Chen et al., 2013, Chistiakova et al., 2015, Fox and Stryker, 2017) to counter the runaway activity of Hebbian plasticity. Integrating our model of structural homeostatic plasticity with Hebbian plasticity could also counter this problem. While most models are difficult to observe in experiments, our model shows an observable sequence of events during learning that could be tested in experiments.

### 3.4 Large-scale formation of memory engrams

To show how well our approach scales, we repeated the experiment with 343 boxes. We assessed that the network behaves correctly for 341 readout neurons firing only with a significantly increased rate when expected. Two readout neurons behaved not as expected. One of the readout neurons fired only at an increased rate for 1,400 steps instead of the entire 2,000 steps of the stimulation of the associated ensemble C1 during the retrieval phase. In contrast, the other readout neuron fired at an increased rate for 500 steps during the retrieval phase when another ensemble C1 was stimulated.

### 3.5 Pattern completion

We want to see if our network can reactivate the learned pattern if we stimulate it only partly. As we see in Fig. S2, the firing rate of the corresponding readout neuron of US increased with more stimulated neurons of C1 except for the change between 0%–5%, 45%–50%, and 80%–85% where the firing rate decreased slightly. We considered a firing rate larger than three times higher than its mean significantly different from its baseline, as 99.7% of the baseline activity lies within this interval (3 *σ* rule (Pukelsheim, 1994)). This was the case for 45% or more stimulated neurons of the ensemble, concluding that at least 150 stimulated neurons are necessary to increase the firing rate of the readout neuron significantly.

Therefore, it is enough to stimulate 45% of the neurons of the ensemble C1 to significantly reactivate the pattern so that the readout neurons fire with a significantly increased firing rate. This shows that we can also perform pattern completion over the engram pairs C1 and US due to our high connectivity within a box.

### 3.6 Forming long-distance connections

As described in Eq. 5, we can control the probability with which neurons connect by adjusting the Gaussian scaling parameter *σ*. We can connect engrams of different boxes when we increase this scaling parameter after we have already formed engrams within a box. To demonstrate this, we increased *σ* and stimulated the ensembles US and C1 from two boxes together and checked in the retrieval phase whether they formed a single, interconnected engram. The firing rates are visualized in Fig. 8. We can see that the readout neurons of both boxes fired once we stimulated them together in step 150,000 and during retrieval when the C1 ensembles of both boxes were stimulated at different times. Thus, we can connect engrams that were not associated before and are distant from each other.

**Figure 8:**
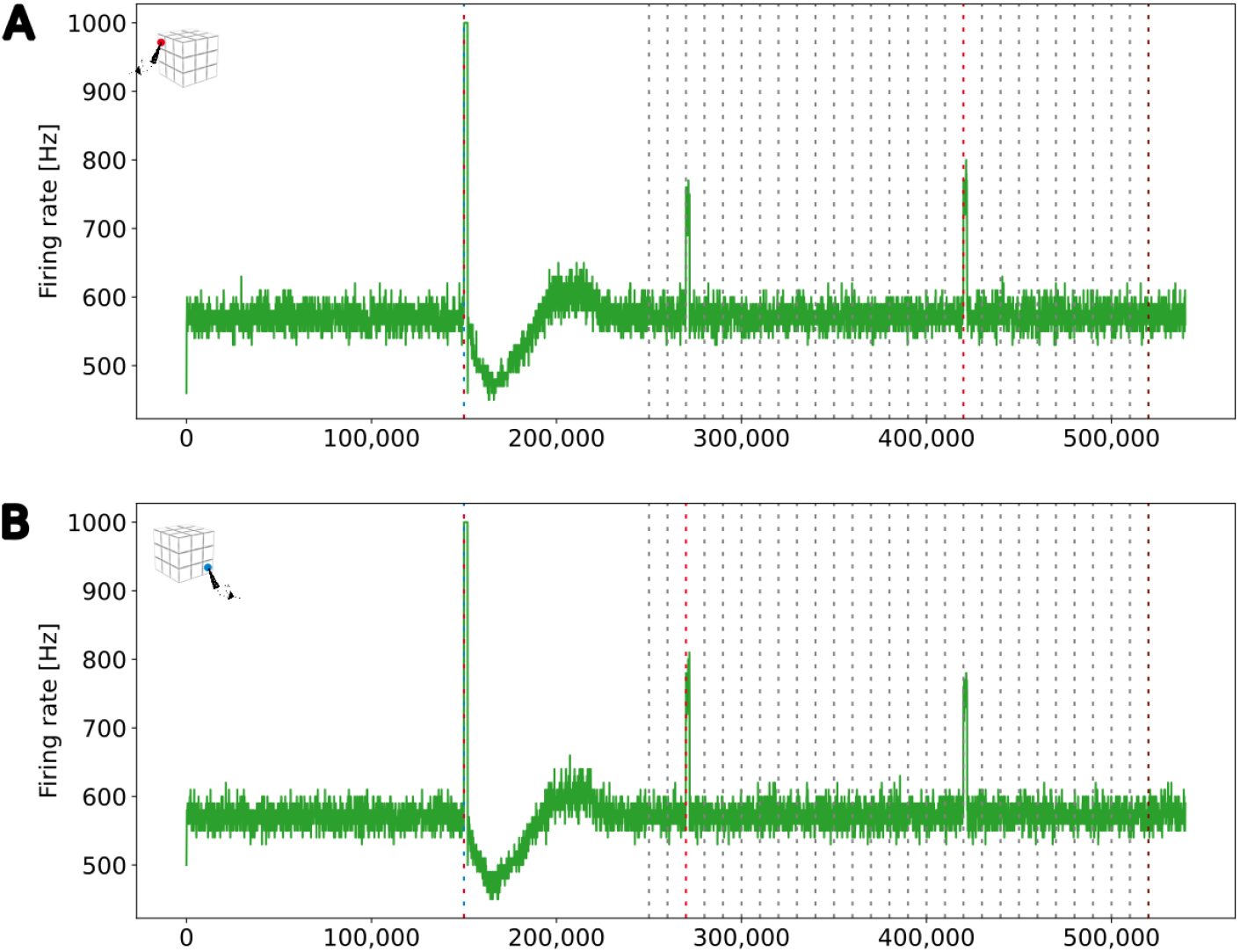
Firing rates of the two readout neurons from the boxes boxes that we stimulated to form long-distance connections. Both neurons fired at a rate of 1 kHz when both ensembles were stimulated together. They also started firing together when only a single ensemble C1 was stimulated, indicating that both boxes had formed a new engram. The blue (red) dashed line mark the stimulation of the ensemble US (C1) of the depending box. **(A)** Firing rate of a read-out neuron located in the upper left box on the top side of the cubic simulation model (see inset A). **(B)** Firing rate of a read-out neuron in the right bottom box at the bottom side of the cubic simulation model (see inset B).

### 3.7 Ablation studies

To see if our model can recover from lesions, we visualize the reaction of a single C1 ensemble to the deletion of all connections from 50% of the neurons of the ensemble in Fig. 9a. Initially, the connectivity from neurons outside US and C1 was by far the strongest. The connectivity from C1 to itself and US to C1 was approximately equal, ranging at about one connection per neuron. We can immediately see the removed connections in step 150,000 as the connectivity from all ensembles decreased drastically. Then, every neuron received less input. Half of the neurons in the ensemble received no synaptic input because we had entirely disconnected them; the other half kept only connections to neurons outside of the ensemble and among each other. As a result, the calcium level started decreasing, and the neurons formed new synapses to rebuild their connections.

**Figure 9:**
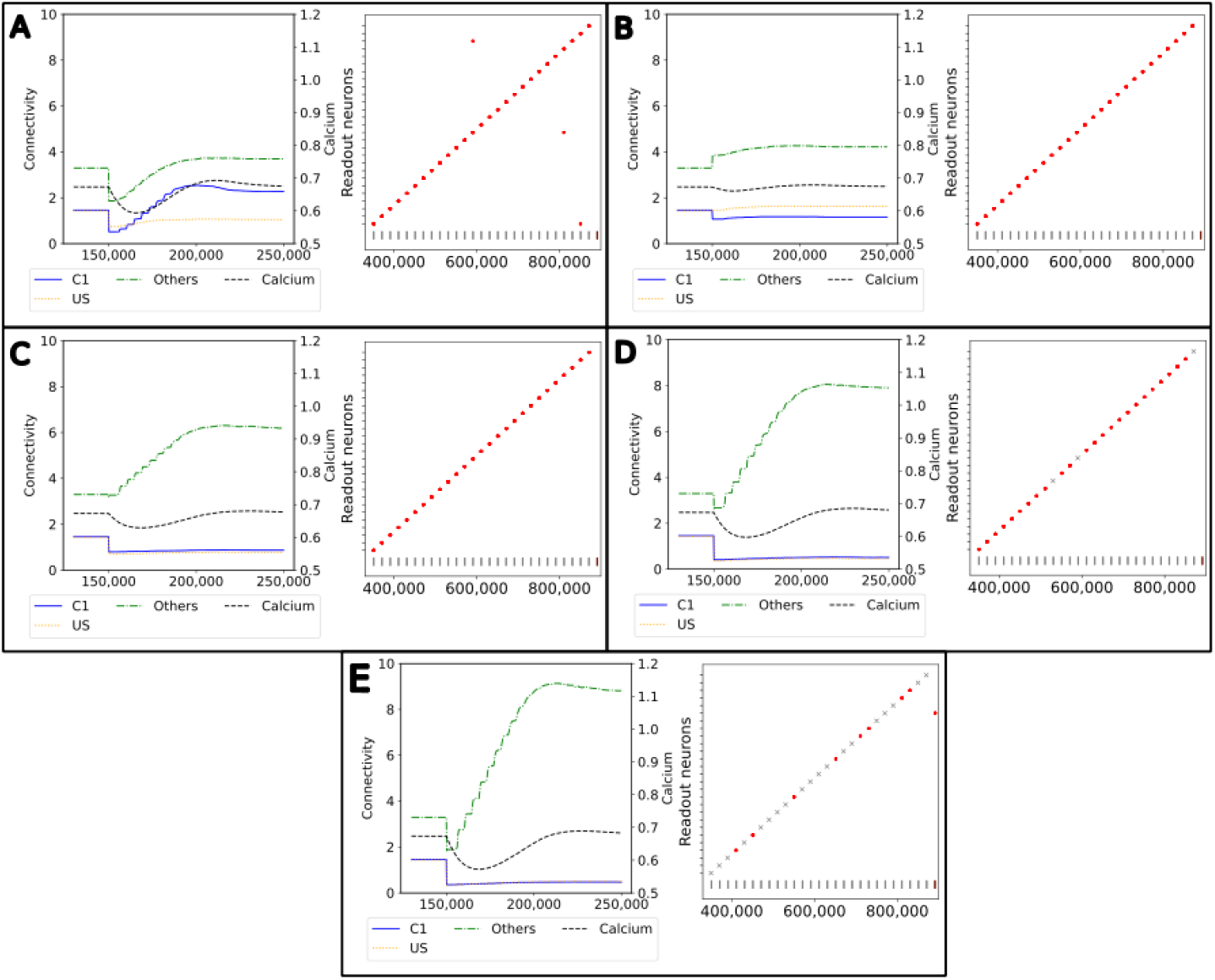
Network response to simulated lesions over time (x-axis). Left: Average input connectivity per neuron for all non-lesioned neurons within a randomly selected example ensemble C1. Input synapses originate from neurons of the same receiving ensemble (blue), from the ensemble US (yellow), or from other ensembles (green) within the same box. The dotted line visualizes the average calcium level of all non-lesioned neurons (right y-axis). A successful functional recovery would require re-formation of input connectivity from ensemble US. Right: Successful functional recovery indicated by red dots meaning increased firing of readout neurons in the retrieval phase as in Figure 2D. Lack of increased firing is marked with a gray cross. The stimulation of single C1 (all C2) ensembles is marked with a vertical gray (brown) bar at the bottom of each diagram. **(A)** Lesioned network loosing the entire connectivity from 50% of the input neurons of ensemble C1. **(B)** Lesioned network in which 50% of the input neurons from the ensemble C1 are completely removed from the network. **(C)** Lesioned network with input connectivity from 30% of randomly selected neurons from the entire box is deleted. **(D)** Lesioned network as in C but with a deletion rate of 50%. **(E)** Lesioned network as in C but with a deletion rate of 70%.

However, the newly grown synapses connected differently than the recently removed ones. The connectivity from US to C1 was approximately only half as strong as before after the network returned to an equilibrium. In exchange, the connectivity from other neurons outside of US and C1 increased slightly. Interestingly, the connectivity within C1 increased to about 2, doubling the connectivity strength within the ensemble. We can explain this behavior with the same effect observed for the learning mechanism with the homeostatic model in Section 3.1. All the neurons from which we removed the connections rebuild their synapses simultaneously so that they start looking for new partners simultaneously. This led to a huge ratio of potential synapse partners within the ensemble. During the retrieval phase at the end, all readout neurons fired at an increased rate at the expected times when we stimulated the C1 of the corresponding box. Nevertheless, the rewiring caused three readout neurons to fire at an increased rate when a C1 from another box was stimulated.

In Fig. 9b, we illustrate the reaction of the network when we permanently apply a lesion to half of the neurons in C1 in step 150,000. A lesion in our model includes removing all connections from and to the lesioned neuron and removing them from the network so that no new connections can be formed. As we completely removed the lesioned neurons from the network, they were no longer considered when calculating the average connectivity after step 150,000. Hence, we see a large decrease in the total connectivity in average connectivity but not in the average connectivity per neuron simply because of the decreased size of the ensemble.

First, the average connectivity shifted immediately after the lesion in step 150,000 because the remaining neurons have slightly different average connectivity. Moreover, the average connectivity from C1 to itself decreased slightly because a fraction of these connections were to lesioned neurons and dropped out. The newly freed synapses can connect instantly with neurons outside US and C1. This leads to a small decrease in the firing rate, which could be observed in the decrease in the average calcium level. Second, the neurons regrew synapses to return to their target activity level. In contrast to our network with only removed connections, mostly the connectivity from neurons outside of US and C1 increased. This is caused by the much smaller fraction of potential synapse partners from neurons within C1, as the lesioned neurons were not looking for new synapse partners. Note that the average connectivity of C1 after the lesions varied due to the randomness involved, as we can see in the shift in the connectivity from other neurons to C1. Other boxes may have different shifts and, therefore, different initial conditions for regrowing their synapses, but all boxes had in common that the connectivity within C1 was not or only slightly strengthened. However, the connectivity from outside of US and C1 was strengthened. Nevertheless, the remaining connectivity from C1 to US was still large enough to enable firing all readout neurons at an increased rate during the retrieval phase.

We visualize the network’s connectivity if we remove all connections from 30% of all neurons within a box regardless of whether they are part of an ensemble in Fig. 9c. The decrease in connectivity in step 150,000 is visible for the connection from US and C1. As the newly freed synapses reconnected immediately to random neurons, we see an increase in the connectivity from neurons outside of US and C1. This increase continued until the neurons were satisfied with their input again. The connectivity from US and C1 remained low as the probability of connecting to these neurons is much smaller due to the small size of the ensembles compared to the rest of the network. However, the remaining connectivity was still sufficient to excite the readout neurons enough during the retrieval phase so that they fired at an increased rate.

If we removed the connections of even more neurons, as visualized in Fig. 9d, the trend was similar but to a larger degree. More connections were removed and restructured mostly to random neurons, leaving only a small number of connections between US and C1. This was visible in the retrieval, and three readout neurons did not fire at an increased rate when we removed 50% of the connections. When we removed all connections from 70% of the neurons (Fig. 9e), the number of connections between US and C1 was close to zero and, therefore, insufficient to trigger an increased firing rate of the readout neurons in 19 out of 27 boxes.

### 3.8 Memory capacity

We will take a closer look at the memory capacity of our model. Fusi (2021) defined the memory capacity of a neural network over the signal-to-noise ratio (SNR), with which memories can be recalled and the number of stored memories. Instead of measuring the neurons’ activity during the memory recall, they observe the changes in the network structure and assume that the changes happening in the network during learning represent a memory. Their model uses synaptic plasticity; therefore, the changed synaptic weights represent memory and the signal. When the network learns more memories, synaptic weights are changed again, reducing the ability to recognize the first memory of the network if it overrides the same synaptic weights. In other words, adding more memories reduces the SNR. This SNR can then be interpreted as an upper bound of the memory capacity—an upper bound because it cannot tell whether the network can recall a memory, only whether its structure is still present in the network, which is necessary but not sufficient. Their work describes that memory is forgotten when its SNR falls below a threshold.

Our model uses only structural plasticity, so we cannot apply the same calculations directly. In our model, each neuron has approximately the same number of synapses *s*_neuron_ in the equilibrium state. We first remove some of the existing synapses of each stimulated neuron and their partners due to the stimulation. Then, the synapses regrow in a structured way so that they encode the newly created memory. The number of retraced and regrown synapses per neuron is roughly the same for each stimulated neuron that we call *s*_engram_ with *s*_engram_ *≤ s*_neuron_. Similar to Fusi (2021), we first calculate the memory signal that can be retrieved directly after creating the memory. Each memory consists of *n*_engram_ number of neurons that we stimulate and recruit. This means that we have 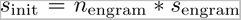 synapses retracted and regrown that represent a single memory initially. As a result, we can express the SNR of the memory in comparison to the noise of the network as proportional to 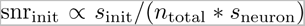. If we add additional memory to the network, we remove and replace *s*_init_ synapses in our network. A single synapse in our memory has the probability of 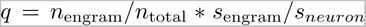 of being removed. The first quotient represents the probability that either the pre- or the postsynaptic neuron belongs to the stimulated neurons, and the second quotient is the probability of a single synapse from a single neuron belonging to the deleted synapses. As a result, the number of synapses still encoding our memory after *t* added memories is proportional to 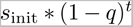 and the SNR is proportional to 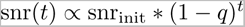.

We visualize the SNR for our model and different parameters in Fig. 10. A higher SNR indicates a better-recallable memory. Our initial SNR depends on the number of neurons in the engram, the number of synapses, and the total number of synapses in the network. We can say that a larger memory—either by a larger number of recruited neurons (Fig. 10d) or a larger number of belonging synapses (Fig. 10b)—increases the initial SNR as we have a lot of synapses (signal) compared to our noise. The larger memory has the disadvantage that we can store fewer memories in our network as synapses are more likely to be overwritten by other synapses. This is visible in the faster fall of the SNR with an increasing number of memories. In contrast, a smaller memory has the disadvantage of an initial small SNR but the advantage of a much slower decreasing SNR as we have more unrelated synapses in our network. This behavior is similar to the plasticity–stability tradeoff described by Fusi (2021) where an increase in synaptic plasticity (e.g., by a large learning rate) allows faster learning of memories but also slower forgetting. In conclusion, our SNR depends on the ratio between the size of a memory compared to the size of the network.

**Figure 10:**
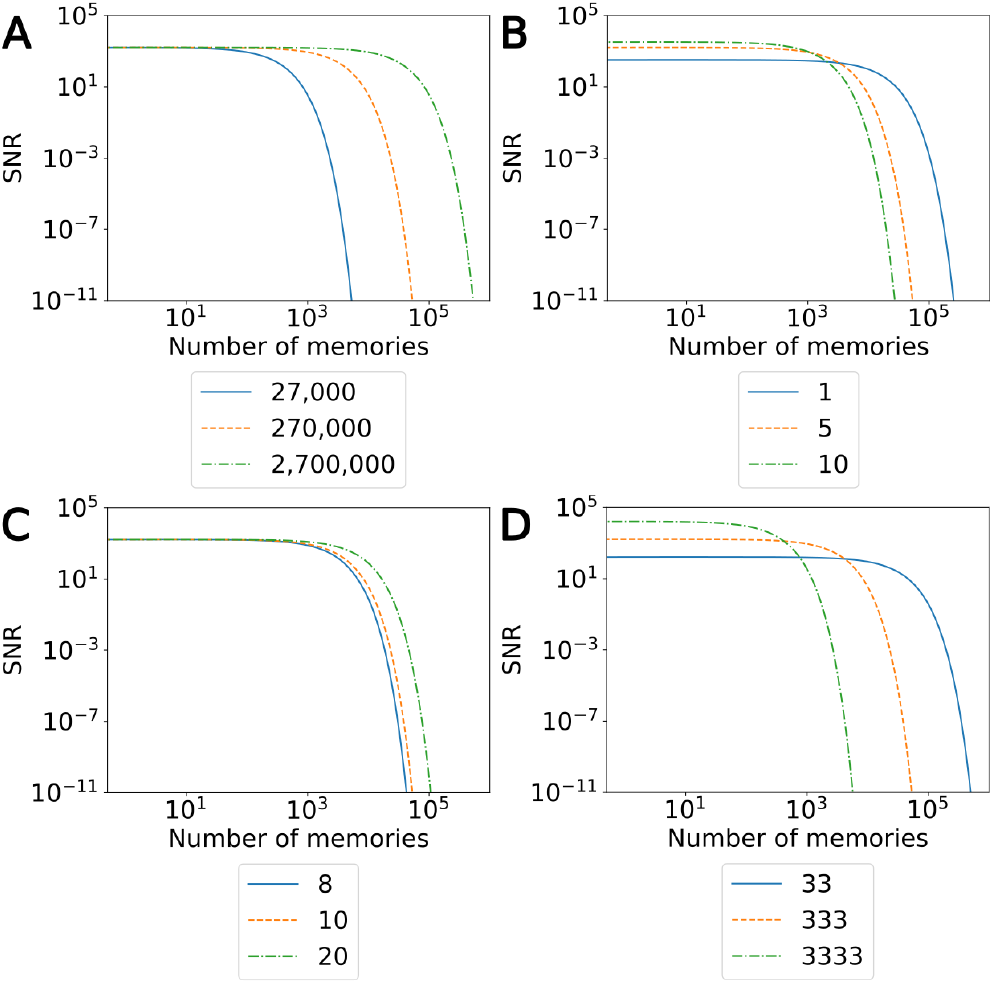
The signal-to-noise ratio (SNR) of our model with varying parameters for an increasing number of stored memories in the network. The orange line visualizes the SNR of the model used in our main experiment with 27 boxes. The blue and green lines visualize the change in the SNR by decreasing (blue) or increasing (green) a single parameter. **(A)** SNR with varying numbers of total neurons in the network. **(B)** SNR with varing number of synapses overwritten per neuron in the engram. **(C)** SNR with varying numbers of synapses per neuron. **(D)** SNR with varying numbers of neurons per engram.

When we talk about large and small memories, we mean the relative size of a memory in ratio to the network size. More precisely, we measure the size of a memory as the total number of synapses a memory occupies. Hence, the larger a memory is, the higher the ratio of synapses out of the total number of synapses in the network. A memory with the same number of synapses is relatively smaller in a network with many neurons than in a small network. Hence, we can increase the SNR of a single memory either by increasing the number of total synapses per memory or by decreasing the number of synapses in the network. Again, the total number of synapses for either a single memory or the entire network can be modified by changing either the number of neurons (Fig. 10a) or the number of synapses per neuron (Fig. 10c). The number of neurons in the network we recruit for a single memory is a direct parameter we can modify in our experiment. On the other hand, we can also indirectly influence the number of synapses as it depends on the calcium level of the neuron. Possible parameters that we can modify to increase the number of synapses per neuron are the target calcium level *η*, the parameters *β* and *τ* that influence the accumulation of the calcium level, or the parameters *ν* and *µ* to influence the growth curve of synaptic elements based on the calcium level. Additionally, if we want to modify the number of synapses retracted and regrown by the stimulation of a neuron, we can modify the stimulation intensity or duration.

Moreover, the information storage capacity could be increased by recruiting neurons to more than one ensemble. This can explain how the human brain can learn many concepts and associations with a limited number of neurons. In our simulations, neurons are mostly in a single memory engram, but forming multiple engrams in a single box with overlapping neurons could significantly increase the number of engrams that can be simulated with the same number of neurons.

## 4 Discussion

We could show that the Model of Structural Plasticity (MSP) (Butz and van Ooyen, 2013) can form multiple and non-interfering memory engrams in a recurrent and sparsely connected network of spiking neurons. This is remarkable as MSP does not aim to form associative memories. With MSP engram formation solely results from the reciprocal interplay of altered activity levels (after stimulation of selected neuronal ensembles and caused by rewired network connectivity) with local homeostatic morphogenetic changes of neurons (forming and deleting of axonal and dendritic elements) and distance-dependent synapse formation and deletion. Hence, MSP emulates associative memory formation with distinct properties that distinguish it from pure Hebbian synaptic plasticity.

### 4.1 Comparison between homeostatic engram formation and synaptic plasticity-driven learning models

In the human brain, synapses are formed only sparsely. With synaptic plasticity alone, only the weights of these synapses can be changed. This can be a problem if learning patterns require strong connectivity between certain neurons, but their connectivity is low (Knoblauch, 2017). Structural plasticity can overcome those suboptimal connectivity patterns of sparsely connected networks by forming new synapses between the required neurons (and reducing cross-talk in neuronal assemblies by pruning synapses). Moreover, this is an important property when the network needs to learn an entirely new input or when the network needs to recover from a partial lesion. Computational models with synaptic plasticity avoid this problem either by all-to-all connectivity (Rolls et al., 2013, Huang and Wei, 2021)—which is computationally demanding on the one hand and not comparable to the brain where connectivity is only sparse on the other hand— or they require the creation of a fixed number of synapses between random neurons (Fiebig and Lansner, 2017, Szatmáary and Izhikevich, 2010, Savin and Triesch, 2014) at the beginning, limiting the ability to form memories between neuron pairs drastically. Our approach overcomes this limitation, allowing the forming of connections between all neurons and influencing the range over which neurons connect by changing our Gaussian probability parameter *σ*.

If we repeat our experiment with only synaptic plasticity with all-to-all connectivity instead of structural plasticity, we would need *n \** (*n −* 1) synapses where *n* is the number of neurons in our network. Our network of 27 boxes consists of 337,500 neurons and 3,009,795 synapses. With all-to-all connectivity, each neuron would need 337,499 synapses compared to our model’s average of 9 synapses, resulting in 113,905,912,500 synapses. If we compare the numbers of our large-scale experiment with 343 boxes and 4,287,500 neurons, we see that while our final network had 38,296,669 synapses, all-to-all connectivity would require about 10^13^ synapses. Even if we connect only a tiny fraction of neuron pairs with synapses, simulating such a large number of synapses is impractical, and the quadratic dependency of the number of synapses on the number of neurons limits the size of the network enormously. In contrast, our network’s synapses grow linearly with the number of neurons, as in our homeostatic model, each neuron aims for the same number of synapses.

An alternative approach could use neuromorphic computing, where specialized hardware circuits are used for the simulations with spiking neuronal networks instead of general-purpose CPUs. This enables optimizations for neuron simulations in the form of power usage, parallelism, and speed. The hardware architectures range from hybrid systems with FPGAs (Park et al., 2016), analog circuits, ASICs (Davies et al., 2018) to massively-parallel supercomputers (Furber et al., 2014). A considerable advantage is the parallel simulation of neurons through direct hardware support, e.g., having a separate circuit for each neuron. The platforms differ in the way that they transmit spikes between the neurons. They can be transmitted over buses (Mortara et al., 1995), grids (Merolla et al., 2007), or special routing networks (Yang et al., 2021b). Synaptic plasticity is also supported by many platforms enabling fast learning (Yang et al., 2021a). These platforms are built biologically motivated, mimicking some aspects of the brain architectures, e.g., a cerebellum network for motor learning (Yang et al., 2021c), dendritic on-line learning (Yang et al., 2023b), and context-dependent learning (Yang et al., 2021a). Based on the platform for context-dependent learning, a real-world application for smart traffic systems was developed (Yang et al., 2023a). Moreover, recent work showed that structural plasticity can be modeled on some of these platforms (Bogdan et al., 2018, Billaudelle et al., 2021). Our approach, which enormously decreases the required number of synapses, could benefit from a neuromorphic platform implementation with an adaptable routing system to further increase its scalability.

### 4.2 Biological implications

We showed that the Model of Structural Plasticity enables the simultaneous formation of multiple memory engrams. We saw clear, distinct spikes during the retrieval phase, indicating that the network learned the relationship between the ensembles US and C1. However, the firing rate of the readout neurons was lower than during the direct stimulation of US with 1 kHz. Moreover, while confirming the general neurophysiological conclusions of Gallinaro et al. (2022), we demonstrated them at a much greater scale—with a network of 4,287,500 neurons and 343 ensemble pairs. And given the favorable scaling properties of the underlying algorithm, we see no obvious limit to further increases.

Our ablation studies showed that the memory engrams are resistant. If a loss of connections affected only neurons within the memory engram, it could recover from it and even strengthen its connections. Even a lesion of 50% of the neurons did not lead to forgetting a memory as the connectivity between the memory ensemble was still high enough. This implies some inherent reserve capacity in cortical connectivity (as currently being discussed for Alzheimer’s disease (Teipel et al., 2016)). If exceeded and connectivity dropped below a critical level, the network would not regrow the connections. As a result, memory engrams may not be recovered.

For this reserve capacity, distant connections may also play a role. Moreover, forming synapses in a distant-dependent manner enables the simulation of ongoing neuron activity in different brain areas without interference. Without this property, all of our ensembles, US and C1, would connect to each other, forming a single large memory engram instead of various smaller ones. Nevertheless, engrams can be largely distributed over the brain brought about by the Gaussian scaling parameter *σ*. An increment of *σ* would be comparable with a phase in which the subject learns a new concept and increases plasticity in the brain. This could be comparable to the release of proteins during learning phases that allow higher plasticity in the brain (Lamprecht and LeDoux, 2004, Poo, 2001). Another possibility is to model different types of neurons with different *σ* to ensure that some neurons can project into other brain areas while others project only locally to their direct neighbors.

### 4.3 Comparison with modular organization of cortical and hippocampal networks

The organization of our 3D model in distinct boxes is comparable to the modular organization of the cortex in columns and hyper-columns (Mountcastle, 1997, Hubel and Wiesel, 1962). One concept is learned in one box very much within a single module of the associative cortices in the brain. However, our model can still learn relationships between more than one box. These memory engrams do not influence the local memory engrams within a box comparable to the human brain, where inactive memories remain unchanged while non-associated memories are active.

Another view on our model is its similarity to the hippocampus, which is an essential area in memory formation (Squire et al., 2004) and shows structural changes during learning (Leuner et al., 2003, Leuner and Gould, 2010, Groussard et al., 2010). Declarative and spatial memory are often associated with the hippocampus (Eichenbaum and Cohen, 2014, Deuker et al., 2016, Eichenbaum, 2017). Spatial memory enables us to navigate and capture the relation of locations. It was originally suspected that the brain consists of a spatial allocentric map that would allow a mapping of locations in an absolute coordinate system (O’Keefe, 1991, Tolman, 1948, O’keefe and Nadel, 1979), but later research discarded this idea (Eichenbaum et al., 1999). Nevertheless, place cells have been found in the hippocampus of humans and rats that are active when the subject is at a certain location (O’Keefe and Dostrovsky, 1971, Hill and Best, 1981, Thompson and Best, 1989, Eichenbaum et al., 1999, Moser et al., 2008). These place cells allow reliable location detection but are not spatially distributed in the hippocampus. However, place cells of the same or nearby location tend to cluster (Eichenbaum et al., 1989), and the distance between these place cells correlates with the real-world locations they represent (O’Keefe and Burgess, 1996). Neurons representing similar concepts, such as similar locations, are often located close to each other. Other examples can be found in the retinotopic organization of the visual cortex (Hubel and Wiesel, 1962, Tusa et al., 1979) and the cerebral cortex (Penfield and Rasmussen, 1950).

This is similar to our model, where engrams are more likely to form locally, leading to a clustering of neurons forming an engram. In our model, one box represents a concept comparable to a place field. With our simulation, we could show that it is possible to form these place cells by simultaneous stimulation comparable to long-term potentation (Teyler and DiScenna, 1987). When rats were placed in a new environment, within minutes, new place cells were activated in the hippocampus and recruited so that they could be recalled weeks later (Rotenberg et al., 1996, Lever et al., 2002). Our model explains this process through the increased activity of the neurons, which leads to a locally limited cluster formation. Only a small cue, such as a perceived environmental one or the recall of the location in the imagination, can reactivate the cluster of place cells (O’Keefe and Conway, 1978). This is similar to our model, where only a partial activation of the unconditioned stimulus (similar to an environmental cue) leads to the reactivation of the conditioned stimulus.

### 4.4 Predictions on the nature of homeostatic engram formation

Our modeling approach predicts that pruning of synapses is an integral part of homeostatic engram formation and does not occur as mere compensatory response to enduring synapse potentiation associated with memory formation (as in Hebbian plasticity with synaptic scaling (Turrigiano, 2008) as synaptic homeostasis rule) but, in fact, precedes and first enables associative synapse formation. Homeostatic engram formation requires a certain spatiotemporal order and availability of vacant axonal and dendritic elements for synapse formation. If vacant synaptic elements are not available in certain ensembles, learning or re-learning of engrams may fail. This is particularly seen in ablation studies and clearly distinguishes homeostatic engram formation from associative Hebbian learning, in which the formation of new engrams merely depends on the coincidence or temporal order of input signals. Homeostatic engram formation can further be boosted by inhibition opening up critical periods for amplified synapse formation (Rinke et al., 2017) and possibly also for engram formation (still to be shown computationally). With this, the present modeling approach provides a number of testable predictions on the nature of homeostatic engram formation.

### 4.5 Conclusions

We believe that our approach lays the foundation for the understanding of more complex memory systems. We could show that the homeostatic engram formation and functional recovery from lesions trigger a neuronal growth mechanism (implemented as homeostatic growth rules) and, hence, may recapitulate neural development. As we have further demonstrated, we can form engrams locally within a box as well as over longer distances by adapting the Gaussian scaling parameter *σ*. Besides the already discussed recruiting of neurons in multiple engrams, we could build associations between multiple engrams locally or distribute them across the network. This would be an important extension because memories in the brain do not exist in a single one-to-one relationship but rather depend on many other memories. Another possibility is to build a memory hierarchy with directed associations where one memory engram activates another but not the other way around. In that case, an abstract concept represented as a memory engram is more strongly associated with another than vice versa. A combination of these extensions would enable the building of a more complex memory system.

## Author Contributions

MK: investigation, methodology, software, writing – original draft, writing – review & editing. FC: methodology, software, supervision, writing – review & editing. MBO: methodology, supervision, writing – review & editing. FW: funding acquisition, methodology, supervision, writing – review & editing.

## Data Availability Statement

The simulator code can be found here: https://github.com/tuda-parallel/RELeARN. Our analysis scripts and data can be found here: https://doi.org/10.5281/zenodo.8287234

## Supplementary Data

**Figure S1:**
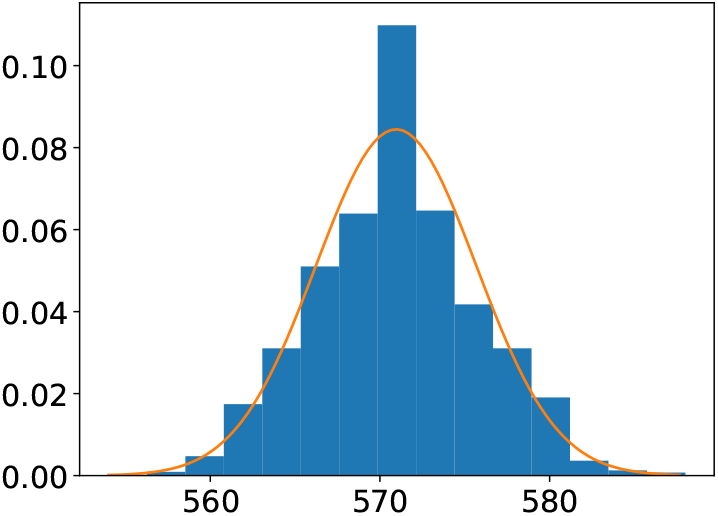
Histogram of the recorded firing rates of the readout neurons (blue bars) and the assumed normal distribution (orange lines) with a mean of 570.95 Hz and a standard deviation of 4.72.

**Figure S2:**
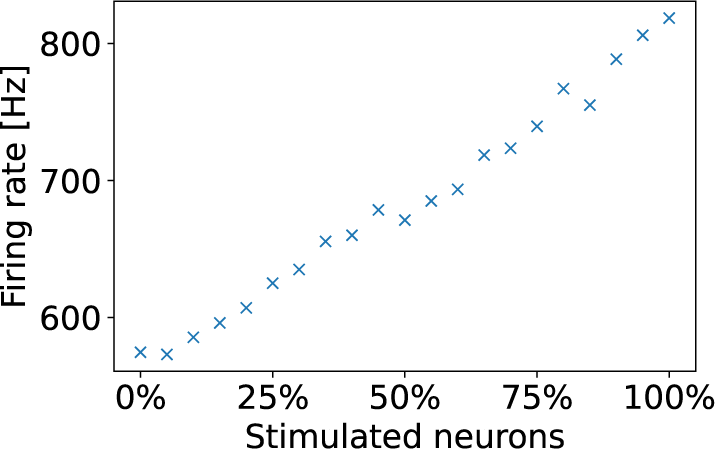
Firing rate during pattern retrieval. The normal firing rate of the readout neuron without any stimulation has a mean of *µ* = 574.5 Hz and a standard deviation *σ* = 29.3. When we stimulated an increasing number of neurons in the ensemble C1, the firing rate of the readout neuron of the associated readout neuron fired at an increasing rate. There are three exception were an increase of the number of stimulated neurons about 5% do not increase the firing rate.

## References

O. Barak and M. Tsodyks. Working models of working memory. Current Opinion in Neurobiology, 25:20–24, 2014. ISSN 09594388. URL 10.1016/j.conb.2013.10.008.

S. J. Barnes and G. T. Finnerty. Sensory experience and cortical rewiring. The Neuroscientist, 16(2):186–198, 2010.

S. Billaudelle, B. Cramer, M. A. Petrovici, K. Schreiber, D. Kappel, J. Schemmel, and K. Meier. Structural plasticity on an accelerated analog neuromorphic hardware system. Neural networks, 133:11–20, 2021.

P. A. Bogdan, A. G. Rowley, O. Rhodes, and S. B. Furber. Structural plasticity on the spinnaker many-core neuromorphic system. Frontiers in Neuroscience, 12:434, 2018.

J. Boyke, J. Driemeyer, C. Gaser, C. Büchel, and A. May. Training-induced brain structure changes in the elderly. Journal of Neuroscience, 28(28):7031–7035, 2008. ISSN 0270-6474. URL https://www.jneurosci.org/content/28/28/7031.

M. Butz and A. van Ooyen. A simple rule for dendritic spine and axonal bouton formation can account for cortical reorganization after focal retinal lesions. PLoS Computational Biology, 9:39–43, 2013. ISSN 1553734X.

M. Butz, F. Wörgötter, and A. V. Ooyen. Activity-dependent structural plasticity. Brain Research Reviews, 60:287–305, 2009. ISSN 0165-0173. URL 10.1016/j.brainresrev.2008.12.023.

P. Caroni, F. Donato, and D. Muller. Structural plasticity upon learning: regulation and functions. Nature Reviews Neuroscience, 13(7):478–490, 2012.

J.-Y. Chen, P. Lonjers, C. Lee, M. Chistiakova, M. Volgushev, and M. Bazhenov. Heterosynaptic plasticity prevents runaway synaptic dynamics. Journal of Neuroscience, 33(40):15915–15929, 2013.

M. Chistiakova, N. M. Bannon, J.-Y. Chen, M. Bazhenov, and M. Volgushev. Homeostatic role of heterosynaptic plasticity: models and experiments. Frontiers in computational neuroscience, 9:89, 2015.

D. B. Chklovskii, B. W. Mel, and K. Svoboda. Cortical rewiring and information storage. Nature, 431, 2004.

F. Czappa, A. Geiß, and F. Wolf. Simulating structural plasticity of the brain more scalable than expected. Journal of Parallel and Distributed Computing, 171:24–27, Jan. 2023. ISSN 0743-7315. URL https://arxiv.org/abs/2210.05267.

I. E. Dammasch. Structural realization of a hebb-type learning rule. In R. M. J. Cotterill, editor, Models of Brain Function, chapter 32, pages 539–552. Cambridge University Press, Cambridge, 1990.

M. Davies, N. Srinivasa, T.-H. Lin, G. Chinya, Y. Cao, S. H. Choday, G. Dimou, P. Joshi, N. Imam, S. Jain, et al. Loihi: A neuromorphic manycore processor with on-chip learning. Ieee Micro, 38(1):82–99, 2018.

L. Deuker, J. L. Bellmund, T. Navarro Schröder, and C. F. Doeller. An event map of memory space in the hippocampus. eLife, 5:e16534, oct 2016. ISSN 2050-084X. URL 10.7554/eLife.16534.

P. E. Downing and M. V. Peelen. The role of occipitotemporal body-selective regions in person perception. Cognitive neuroscience, 2(3-4):186–203, 2011.

H. Eichenbaum. On the integration of space, time, and memory. Neuron, 95(5): 1007–1018, 2017. ISSN 0896-6273. URL https://www.sciencedirect.com/science/article/pii/S0896627317305603.

H. Eichenbaum and N. J. Cohen. Can we reconcile the declarative memory and spatial navigation views on hippocampal function? Neuron, 83(4):764–770, 2014.

H. Eichenbaum, S. I. Wiener, M. Shapiro, and N. Cohen. The organization of spatial coding in the hippocampus: a study of neural ensemble activity. Journal of Neuroscience, 9(8):2764–2775, 1989.

H. Eichenbaum, P. Dudchenko, E. Wood, M. Shapiro, and H. Tanila. The hippocampus, memory, and place cells: is it spatial memory or a memory space? Neuron, 23(2):209–226, 1999.

F. Fiebig and A. Lansner. A spiking working memory model based on hebbian short-term potentiation. Journal of Neuroscience, 37(1):83–96, 2017. ISSN 0270-6474. URL https://www.jneurosci.org/content/37/1/83.

K. Fox and M. Stryker. Integrating hebbian and homeostatic plasticity: introduction, 2017.

S. B. Furber, F. Galluppi, S. Temple, and L. A. Plana. The spinnaker project. Proceedings of the IEEE, 102(5):652–665, 2014.

S. Fusi. Memory capacity of neural network models. arXiv preprint arXiv:2108.07839, 2021.

J. V. Gallinaro, N. Gašparović, and S. Rotter. Homeostatic control of synaptic rewiring in recurrent networks induces the formation of stable memory engrams. PLoS Computational Biology, 18, 2 2022. ISSN 15537358.

M. Groussard, R. La Joie, G. Rauchs, B. Landeau, G. Chetelat, F. Viader, B. Desgranges, F. Eustache, and H. Platel. When music and long-term memory interact: effects of musical expertise on functional and structural plasticity in the hippocampus. PLoS One, 5(10):e13225, 2010.

N. Hadjikhani, A. K. Liu, A. M. Dale, P. Cavanagh, and R. B. Tootell. Retinotopy and color sensitivity in human visual cortical area v8. Nature neuroscience, 1 (3):235–241, 1998.

A. Hayashi-Takagi, S. Yagishita, M. Nakamura, F. Shirai, Y. I. Wu, A. L. Loshbaugh, B. Kuhlman, K. M. Hahn, and H. Kasai. Labelling and optical erasure of synaptic memory traces in the motor cortex. Nature, 525(7569): 333–338, 2015.

A. J. Hill and P. J. Best. Effects of deafness and blindness on the spatial correlates of hippocampal unit activity in the rat. Experimental neurology, 74 (1):204–217, 1981.

A. Holtmaat and P. Caroni. Functional and structural underpinnings of neuronal assembly formation in learning. Nature Neuroscience, 19:1553–1562, 2016. ISSN 15461726. URL 10.1038/nn.4418.

A. Holtmaat and K. Svoboda. Experience-dependent structural synaptic plasticity in the mammalian brain. Nature Reviews Neuroscience, 10:647–658, 2009. ISSN 1471003X.

Q.-S. Huang and H. Wei. A computational model of working memory based on spike-timing-dependent plasticity. Frontiers in Computational Neuroscience, 15:630999, 2021.

D. H. Hubel and T. N. Wiesel. Receptive fields, binocular interaction and functional architecture in the cat’s visual cortex. The Journal of physiology, 160(1):106, 1962.

E. Izhikevich. Which model to use for cortical spiking neurons? IEEE Transactions on Neural Networks, 15(5):1063–1070, 2004.

N. Kalisman, G. Silberberg, and H. Markram. The neocortical microcircuit as a tabula rasa. Proceedings of the National Academy of Sciences, 102(3):880–885, 2005. URL https://www.pnas.org/doi/abs/10.1073/pnas.0407088102.

A. Knoblauch. Chapter 17 - impact of structural plasticity on memory formation and decline. In A. van Ooyen and M. Butz-Ostendorf, editors, The Rewiring Brain, pages 361–386. Academic Press, San Diego, 2017. ISBN 978-0-12-803784-3. doi: 10.1016/B978-0-12-803784-3.00017-2. URL https://www.sciencedirect.com/science/article/pii/B9780128037843000172.

R. Lamprecht and J. LeDoux. Structural plasticity and memory. Nature Reviews Neuroscience, 5(1):45–54, 2004.

B. Leuner and E. Gould. Structural plasticity and hippocampal function. Annual review of psychology, 61:111–140, 2010.

B. Leuner, J. Falduto, and T. J. Shors. Associative memory formation increases the observation of dendritic spines in the hippocampus. Journal of Neuroscience, 23(2):659–665, 2003.

C. Lever, T. Wills, F. Cacucci, N. Burgess, and J. O’Keefe. Long-term plasticity in hippocampal place-cell representation of environmental geometry. Nature, 416(6876):90–94, 2002.

M. F. López-Aranda, J. F. López-Téllez, I. Navarro-Lobato, M. Masmudi-Martín, A. Gutiérrez, and Z. U. Khan. Role of layer 6 of v2 visual cortex in object-recognition memory. Science, 325(5936):87–89, 2009.

F. J. Massey Jr. The kolmogorov-smirnov test for goodness of fit. Journal of the American statistical Association, 46(253):68–78, 1951.

A. May. Experience-dependent structural plasticity in the adult human brain. Trends in Cognitive Sciences, 15:475–482, 2011. ISSN 13646613. URL 10.1016/j.tics.2011.08.002.

P. A. Merolla, J. V. Arthur, B. E. Shi, and K. A. Boahen. Expandable networks for neuromorphic chips. IEEE Transactions on Circuits and Systems I: Regular Papers, 54(2):301–311, 2007.

A. Mizrahi. Dendritic development and plasticity of adult-born neurons in the mouse olfactory bulb. Nature neuroscience, 10(4):444–452, 2007.

A. Mortara, E. A. Vittoz, and P. Venier. A communication scheme for analog vlsi perceptive systems. IEEE Journal of Solid-State Circuits, 30(6):660–669, 1995.

E. I. Moser, E. Kropff, and M.-B. Moser. Place cells, grid cells, and the brain’s spatial representation system. Annu. Rev. Neurosci., 31:69–89, 2008.

V. B. Mountcastle. The columnar organization of the neocortex. Brain: a journal of neurology, 120(4):701–722, 1997.

J. O’Keefe. An allocentric spatial model for the hippocampal cognitive map. Hippocampus, 1(3):230–235, 1991.

J. O’Keefe and N. Burgess. Geometric determinants of the place fields of hippocampal neurons. Nature, 381(6581):425–428, 1996.

J. O’Keefe and D. H. Conway. Hippocampal place units in the freely moving rat: why they fire where they fire. Experimental brain research, 31:573–590, 1978.

J. O’Keefe and J. Dostrovsky. The hippocampus as a spatial map: preliminary evidence from unit activity in the freely-moving rat. Brain research, 1971.

J. O’keefe and L. Nadel. Précis of o’keefe & nadel’s the hippocampus as a cognitive map. Behavioral and Brain Sciences, 2(4):487–494, 1979.

J. Park, T. Yu, S. Joshi, C. Maier, and G. Cauwenberghs. Hierarchical address event routing for reconfigurable large-scale neuromorphic systems. IEEE transactions on neural networks and learning systems, 28(10):2408–2422, 2016.

W. Penfield and T. Rasmussen. The cerebral cortex of man; a clinical study of localization of function. Journal of the American Medical Association, 144 (16):1412–1412, 12 1950. ISSN 0002-9955. URL 10.1001/jama.1950.02920160086033.

M.-m. Poo. Neurotrophins as synaptic modulators. Nature reviews neuroscience, 2(1):24–32, 2001.

F. Pukelsheim. The three sigma rule. The American Statistician, 48(2):88–91, 1994.

R. A. Reale and T. J. Imig. Tonotopic organization in auditory cortex of the cat. Journal of Comparative Neurology, 192(2):265–291, 1980.

S. Rinke, M. Naveau, F. Wolf, and M. Butz-Ostendorf. Chapter 8 - critical periods emerge from homeostatic structural plasticity in a full-scale model of the developing cortical column. In A. van Ooyen and M. Butz-Ostendorf, editors, The Rewiring Brain, pages 177–201. Academic Press, San Diego, 2017. ISBN 978-0-12-803784-3. doi: 10.1016/B978-0-12-803784-3.00008-1. URL https://www.sciencedirect.com/science/article/pii/B9780128037843000081.

S. Rinke, M. Butz-Ostendorf, M. A. Hermanns, M. Naveau, and F. Wolf. A scalable algorithm for simulating the structural plasticity of the brain. Journal of Parallel and Distributed Computing, 120:251–266, 2018. ISSN 07437315. URL 10.1016/j.jpdc.2017.11.019.

E. T. Rolls, L. Dempere-Marco, and G. Deco. Holding multiple items in short term memory: a neural mechanism. PloS one, 8(4):e61078, 2013.

A. Rotenberg, M. Mayford, R. D. Hawkins, E. R. Kandel, and R. U. Muller. Mice expressing activated camkii lack low frequency ltp and do not form stable place cells in the ca1 region of the hippocampus. Cell, 87(7):1351–1361, 1996.

J. Ruben, J. Schwiemann, M. Deuchert, R. Meyer, T. Krause, G. Curio, K. Villringer, R. Kurth, and A. Villringer. Somatotopic organization of human secondary somatosensory cortex. Cerebral cortex, 11(5):463–473, 2001.

C. Savin and J. Triesch. Emergence of task-dependent representations in working memory circuits. Frontiers in computational neuroscience, 8:57, 2014.

L. R. Squire, C. E. Stark, and R. E. Clark. The medial temporal lobe. Annual Review of Neuroscience, 27(1):279–306, 2004.

A. Stepanyants, P. R. Hof, and D. B. Chklovskii. Geometry and structural plasticity of synaptic connectivity. Neuron, 34(2):275–288, 2002. ISSN 0896-6273. URL https://www.sciencedirect.com/science/article/pii/S0896627302006529.

B. Szatmáry and E. M. Izhikevich. Spike-timing theory of working memory. PLoS computational biology, 6(8):e1000879, 2010.

S. Teipel, M. J. Grothe, J. Zhou, J. Sepulcre, M. Dyrba, C. Sorg, and C. Babiloni. Measuring cortical connectivity in alzheimer’s disease as a brain neural network pathology: toward clinical applications. Journal of the International Neuropsychological Society, 22(2):138–163, 2016.

T. J. Teyler and P. DiScenna. Long-term potentiation. Annual review of neuroscience, 10(1):131–161, 1987.

L. Thompson and P. Best. Place cells and silent cells in the hippocampus of freely-behaving rats. Journal of Neuroscience, 9(7):2382–2390, 1989.

E. C. Tolman. Cognitive maps in rats and men. Psychological review, 55(4):189, 1948.

J. T. Trachtenberg, B. E. Chen, G. W. Knott, G. Feng, J. R. Sanes, E. Welker, and K. Svoboda. Long-term in vivo imaging of experience-dependent synaptic plasticity in adult cortex. Nature, 420(6917):788–794, 2002.

G. G. Turrigiano. The self-tuning neuron: synaptic scaling of excitatory synapses. Cell, 135(3):422–435, 2008.

R. Tusa, A. Rosenquist, and L. Palmer. Retinotopic organization of areas 18 and 19 in the cat. Journal of Comparative Neurology, 185(4):657–678, 1979.

T. Xu, X. Yu, A. J. Perlik, W. F. Tobin, J. A. Zweig, K. Tennant, T. Jones, and Y. Zuo. Rapid formation and selective stabilization of synapses for enduring motor memories. Nature, 462(7275):915–919, 2009.

S. Yang, J. Wang, B. Deng, M. R. Azghadi, and B. Linares-Barranco. Neuromorphic context-dependent learning framework with fault-tolerant spike routing. IEEE Transactions on Neural Networks and Learning Systems, 33 (12):7126–7140, 2021a.

S. Yang, J. Wang, X. Hao, H. Li, X. Wei, B. Deng, and K. A. Loparo. Bicoss: toward large-scale cognition brain with multigranular neuromorphic architecture. IEEE Transactions on Neural Networks and Learning Systems, 33(7): 2801–2815, 2021b.

S. Yang, J. Wang, N. Zhang, B. Deng, Y. Pang, and M. R. Azghadi. Cerebellumorphic: large-scale neuromorphic model and architecture for supervised motor learning. IEEE Transactions on Neural Networks and Learning Systems, 33(9):4398–4412, 2021c.

S. Yang, J. Tan, T. Lei, and B. Linares-Barranco. Smart traffic navigation system for fault-tolerant edge computing of internet of vehicle in intelligent transportation gateway. IEEE Transactions on Intelligent Transportation Systems, 2023a.

S. Yang, H. Wang, Y. Pang, M. R. Azghadi, and B. Linares-Barranco. Nadol: Neuromorphic architecture for spike-driven online learning by dendrites. IEEE Transactions on Biomedical Circuits and Systems, 2023b.

